# RepeatOBserver: tandem repeat visualization and centromere detection

**DOI:** 10.1101/2023.12.30.573697

**Authors:** Cassandra Elphinstone, Rob Elphinstone, Marco Todesco, Loren Rieseberg

## Abstract

Tandem repeats can play an important role in centromere structure, subtelomeric regions, DNA methylation, recombination, and the regulation of gene activity. There is a growing need for bioinformatics tools that can visualize and explore chromosome-scale repeats. Here we present RepeatOBserver, a new tool for visualizing tandem repeats and clustered transposable elements and for identifying potential natural centromere locations, using a Fourier transform of DNA walks: https://github.com/celphin/RepeatOBserverV1. RepeatOBserver can identify a broad range of repeats (3-20, 000bp long) in genome assemblies without any *a priori* knowledge of repeat sequences or the need for optimizing parameters. RepeatOBserver allows for easy visualization of the positions of both perfect and imperfect repeating sequences across each chromosome. We use RepeatOBserver to compare DNA walks, repeat patterns and centromere positions across genome assemblies in a wide range of well-studied species (e.g., human, mouse-ear cress), crops, and non-model organisms (e.g., fern, yew). Analyzing 107 chromosomes with known centromere positions, we find that centromeres consistently occur in regions that have the least diversity in repeat types (i.e. one or a few repeated sequences are present in very high numbers). Taking advantage of this information, we use a genomic Shannon diversity index to predict centromere locations in several other chromosome-scale genome assemblies. The Fourier spectra produced by RepeatOBserver can help visualize historic centromere positions, potential neocentromeres, retrotransposon clusters and gene copy variation. Identification of patterns of split and inverted tandem repeats at inversion boundaries suggests that at least some chromosomal inversions or misassemblies can be predicted with RepeatOBserver. RepeatOBserver is therefore a flexible tool for comprehensive characterization of tandem repeat patterns that can be used to visualize and identify a variety of regions of interest in genome assemblies.

## Introduction

The increasing availability of chromosome-scale genome assemblies for a wide range of non-model species, as well as fully haplotype-resolved assemblies and pangenomes for well-studied species (e.g. *Homo sapiens, Solanum* spp., *Sorghum bicolor, Gallus gallus domesticus*) (Li *et al*., 2022; Gong *et al*., 2023; Liao *et al*., 2023; Li *et al*., 2023; Ruperao *et al*., 2021; Wang *et al*., 2021, Wlodzimierz, Rabanal, *et al*., 2023), is opening a world of possibilities for looking at non-genic regions (including tandem repeats, centromeres) and genome structural evolution. Along with improving long read and phasing sequencing technologies, bioinformatics tools are needed to better visualize genome-wide patterns of sequence and structural variation in these genome assemblies (Miga, 2020). For example, being able to predict the potential locations of centromeres without the need for direct experimental detection (e.g., fluorescent in-situ hybridization, FISH), accurate genome annotations or information on DNA methylation patterns would be extremely useful for understanding genome structure and evolution (Hartley and O’Neill, 2019; Wlodzimierz, Hong, et al., 2023). Likewise, visualizing tandem repeat patterns throughout the genome would facilitate identification of other regions of potential interest (e.g. inversions, neocentromeres and gene copy number variation).

### Centromeres

Centromeres play a crucial role in chromosome segregation during cell division and knowing their locations helps predict recombination rates across chromosomes. Centromeres are difficult to reliably identify since they undergo rapid evolution due to centromeric drive (Henikoff *et al.,* 2001; Kursel and Malik, 2018; Talbert and Henikoff, 2022; Bracewell *et al*., 2019; Lampson and Black, 2017; Fishman and Saunders, 2008, Melters *et al*., 2012; Wlodzimierz, Rabanal, *et al*., 2023). Centromere drive occurs when new centromeres act selfishly and trigger unequal segregation during female meiosis making one centromere and its chromosome more likely than the other old homologous chromosome to be inherited. Centromeres are made of a combination of epigenetic (e.g. histone methylation) and genomic (often AT-rich, satellite DNA) features (Saha, 2019; Naish *et al*., 2021; Gent *et al*., 2012; Talbert and Henikoff, 2020). Chromosomes can have one centromere (point, regional, or satellite), two centromeres (dicentric), or even centromeres distributed across the whole chromosome (holocentric) (Scott and Sullivan, 2014; Melters *et al*., 2012; Talbert and Henikoff, 2020). *De novo* or neo-centromeres can form due to a shifted epigenetic signature, often to a position without the same underlying sequence; in humans, this can lead to some types of cancer and other extreme phenotypes (Amor and Choo, 2002). Typically, centromeres are identified in the lab using FISH probes, CHIPseq, CENP-A footprinting, or chromatin fiber analysis (Saha, 2019; Nietzel *et al*., 2001). Tandem repeats have been shown to often occur at the locations of centromeres (Melters *et al*., 2013; Hartley and O’Neill, 2019) and it has even been suggested that the number of repeats at the centromere strengthens centromere binding during meiosis causing centromeric drive (Talbert and Henikoff, 2020). However, in many newly formed or artificial centromeres, there are no large tandem repeat arrays, implying that these repeats form later in centromere development (Hartley and O’Neill, 2019). In some domesticated species with recent centromere shifts, such as *Zea mays*, *Gallus gallus domesticus* and *Oryza sativa*, their centromeres lack tandem repeats but often contain retrotransposons (Talbert and Henikoff, 2020). Past studies have explored and classified centromeric repeat sequence similarities across many species (Melters *et al*., 2013; Talbert and Henikoff, 2020).

Centromere locations have only been predicted in a few species using bioinformatics. One approach used wavelet analyses of gene densities and DNA methylation patterns to detect centromeres in *Populus trichocarpa* (Weighill *et al*., 2019). However, this analysis requires methylation data and an accurate genome annotation which is not available for all non-model genome assemblies. Another proposed method uses patterns in HiC maps to predict centromere locations (Varoquaux et al., 2015). Other studies have shown links between CG isochores (Costantini *et al*., 2006; Zhang and Zhang, 2004; Jabbari and Bernardi, 2017; Bernardi, 2017; Woody *et al.,* 2012; Lynch *et al*., 2010) and centromere positions in some specific species. A recent method identifies many possible centromere locations using repetitive DNA (Lin *et al.,* 2023). Overall, these programs often require complex datasets (e.g. methylation, HiC, or gene/TE annotations) and typically lack simple chromosome-scale visualization of repeat structure.

Other bioinformatics tools annotate tandem repeat and/or transposable elements for model reference genomes and their centromeres (TRASH (Wlodzimierz, Hong, *et al.,* 2023), CentromereArchitect (Dvorkina *et al*., 2021), HiCAT (Gao *et al*., 2023) and Repeat Modeler (Flynn *et al*., 2020)). These programs do not predict centromere positions but annotate repeat structures within known centromeres. Most rely on knowing the tandem (centromeric) repeats for a given species, limiting these analyses to model species with known repeats (HiCAT and CentromereArchitect). One exception is TRASH which does not need the sequence of Higher Order Repeats (HORs) to be known but uses a kmer technique that requires the tandem repeats it detects to be close to exact matches, preventing annotation of some imperfect repeats (Wlodzimierz, Hong, *et al.,* 2023).

### Tandem repeats

Tandem repeats appear not only in centromeres but also play an important role in determining telomeres and subtelomeric regions, and patterns of recombination and inversions. They can also contribute to genetic disorders by affecting gene regulation (Horton *et al*., 2023). Many assembled chromosomes include nearly perfect tandem telomeric repeats (TTTAGGG in plants (Richards and Ausubel, 1988), TTGGGG in protists and TTAGGG in animals (Greider and Blackburn, 1985; Kilburn *et al*., 2001)) that can span a few thousand base pairs. Adjacent to these telomeric repeats, there are often subtelomeric repeats which cover a larger portion of the chromosome (sometimes a few Mbp) (Kwapisz and Morillon, 2020), which are often semi-regular and can be conserved between different species. Coding and noncoding RNA are transcribed from these subtelomeric regions and are believed to help with the maintenance of the telomere (Kwapisz and Morillon, 2020).

Although most repetitive DNA sequences (e.g. long terminal repeat retrotransposons) tend to accumulate in regions of chromosomes that have lower rates of recombination (Tiley and Burleigh, 2015), some tandem repeats (specifically GT-rich ones) have been suggested to be recombination hotspots (Majewski and Ott, 2000; Bzymek and Lovett, 2001). During DNA replication, if there is slippage within a tandemly repeated region, this can change the number of repeats. Similarly, non-homologus recombination between repeated sequences can result in chromosomal inversions (Flores *et al*., 2007), and inversions have been found to be occasionally bracketed by split tandem repeats (Hirabayashi and Owens, 2022). Finally, increases in the abundance of short tandem repeats (STR), in particular trinucleotide repeats, are associated with some genetic disorders in both *Homo sapiens* and *Arabidopsis* (Sureshkumar *et al*., 2009). Gene regulation is also affected by STRs that can bind transcription factors (Horton *et al*., 2023). Overall, both short and long tandem repeats play an important role in many aspects of genome organization and function.

Several heuristic programs have been developed to identify tandem repeats of a wide range of lengths in whole-genome assemblies. One of the first, and most frequently used, is Tandem Repeat Finder, which uses Bernoulli trials of kmers to statistically detect candidate tandem repeats (Benson, 1999). This program requires the user to set input parameters that allow for varying amounts of mutation or error in the repeats they can identify. More recently, other programs such as mreps (Kolpakov *et al*., 2003), Spectral Repeat Finder (SRF) (Sharma *et al*., 2004), ATRHunter (Wexler *et al*., 2005), TRStalker (Pellegrini *et al*., 2010), and Tide Hunter (Gao *et al*., 2019) have been designed to better search for fuzzy or imperfect tandem repeats. K-Seek is a tool that has been used in *Drosophila* genomes to find common tandem repeats in Illumina short reads (Wei et al., 2014). There are also databases of common tandem repeats that can be used to compare between species and aid in annotation (Navajas-Pérez and Paterson, 2009). None of these tandem repeat detection programs determine centromere locations based on these repeats and most programs require the user to adjust input parameters to detect all types of repeats.

### RepeatOBserver

We developed a new tandem repeat visualization tool, RepeatOBserver, to explore chromosome-wide patterns of tandem repeats and predict centromere positions in model and non-model species. Starting from only a fasta file for a given chromosome, RepeatOBserver produces Fourier spectra showing locations of tandem repeats (including their length and how perfectly they repeat). Fourier transforms and wavelet analyses have previously been used to study DNA, including exploring DNA periodicity (Elloumi *et al*., 2012; Nagai *et al*., 2001, 2020), DNA palindromes (Qi *et al*., 2012), sequence comparison (MAFFT) (Katoh *et al*., 2002), sequence evolution (Machado, 2013), exon/intron identification (Haimovich *et al*., 2006), visualization of regular features (Dodin *et al*., 2000) and even tandem repeats (Sharma *et al*., 2004; Brodzik, 2007; Buchner and Janjarasjitt, 2003; Yadav *et al*., 2022). DNA sequence variations in Fourier transforms have previously been shown to have biological importance (Haimovich *et al*., 2006) and are known to show periodicity and tandem repeats (Elloumi *et al*., 2012; Sharma *et al*., 2004; Brodzik, 2007; Buchner and Janjarasjitt, 2003; Yadav *et al*., 2022).

Here, we apply Fourier transforms to chromosome scale AT or CG DNA walks to provide an accessible tool for viewing tandem repeat patterns and predicting centromere locations in chromosome-level genome assemblies based on repeat abundance and diversity, without relying on data on epigenetic markers (such as DNA methylation) or gene annotations. RepeatOBserver also allows for the easy visualization of positions of potential neocentromeres, gene copy number variation, repeats around inversions and other short and long tandem repeat patterns.

The advantages of RepeatOBserver when compared to other tandem repeat detection software (Supplementary Table S0) are: (1) The ability to predict centromere positions in many chromosomes based on their repeat structure alone; (2) facile detection of tandem repeats without any prior knowledge; (3) ability to detect a wide range of repeat lengths (3-10 000 bp); (4) capacity to identify extremely imperfect tandem repeats (including retrotransposons), since it does not rely on kmers; (5) easy visualization of whole chromosomes; and (6) lack of a need for optimizing any input parameters.

## Methods

### R package

The R package RepeatOBserver can be found on github: https://github.com/celphin/RepeatOBserverV1 and can be run locally (1 cpu) or with multiple cpu locally or on a server. Scripts for running all the code described in this paper are included in the repository.

### 2D DNA walks

The R package RepeatOBserver first converts the chromosome fasta file input into a numerical DNA walk. Beginning at (0, 0) and stepping through each nucleotide in the chromosome, the 2D DNA walk moves horizontally along the x axis by +1 for each C and by -1 for each G in the genome (Figure 1A). Similarly, the walk moves vertically along the y axis by +1 for A and -1 for T. Plots of this numerical DNA walk allow the overall chromosome structure and large-scale tandem repeats to be viewed (Berger *et al*., 2004). Dominance in any nucleotide or set of repeating nucleotides can be viewed as a slow movement along a vector in the 2D walk.

**Figure 1:**
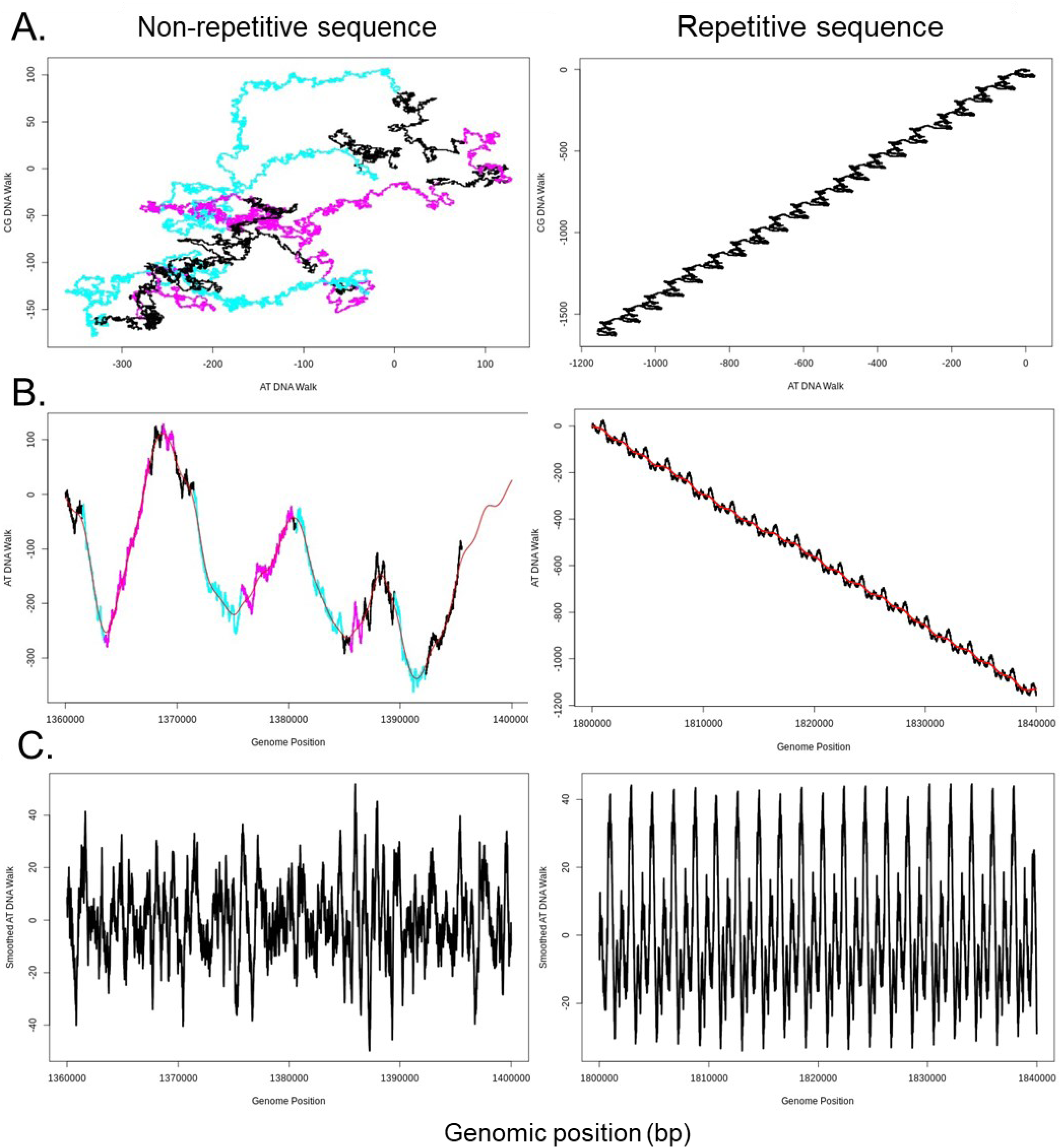
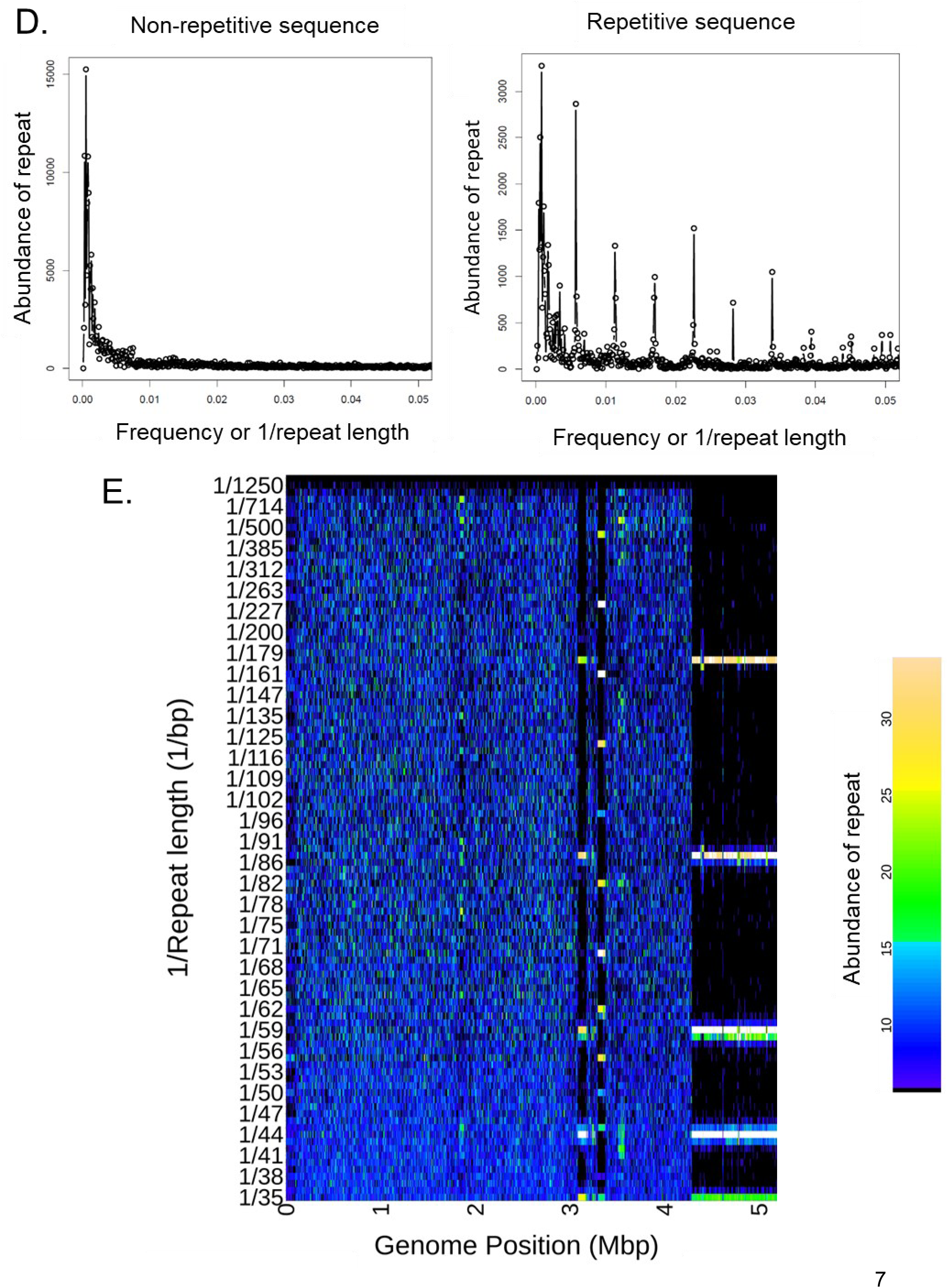
Example 40 kbp sequences from the *Arabidopsis thaliana* genome (COLCEN) (Naish et al., 2021) chromosome 4 showing a non-repetitive section of the genome (left) and a repetitive sequence (right). (A) Plot of a 2D DNA walk with the Araport11 annotation showing forward genes plotted in blue and reverse genes plotted in pink (Cheng *et al*., 2017). (B) 1D AT DNA walk with a smoothing spline (red line), again forward genes plotted in blue and reverse genes plotted in pink. (C) smoothed sequence prior to the Fourier Transform (D) Fast Fourier Transform output showing the relative abundance of the different frequencies (with frequency represented by 1/repeat length in bp). The largest repeat length (smallest frequency) is the fundamental and the length of the repeating DNA sequence; the harmonics are also shown and describe the complexity of the repeat. (E) Heat map showing all the 5000bp spectra joined together (to show a 5Mbp region), with the x axis showing genome position, the y axis plotting the frequency (1/repeat length) and the colour showing the abundance of the repeat at that frequency in that position of the genome.

### 1D DNA walks

One dimensional DNA walks are plotted and used for the Fourier transform input. A one-dimensional AT walk starts at (0, 0) and steps along the x-axis for each base pair in the genome sequence. On the y-axis, each A in the sequence increments the plot by +1, each T by -1 and both C and G increment by 0 (Figure 1B). Similarly, the CG walk starts at (0, 0) and steps along the x-axis for each base pair in the genome sequence. On the y-axis, each C in the sequence increments the plot by +1, each G by -1 and both A and T increment by 0. When gene annotations are plotted on the 1D DNA walks, forward genes (blue) most often appear in the AT walk as negative slopes and reverse genes (pink) often as positive slopes. This pattern likely results from a higher frequency of codons enriched in T and explains some of the DNA walk movement outside of the repetitive regions.

### Smoothing spline

Over large regions of the genome there can be a trend in base pairs that is not directly related to the shorter repeats of interest. This large-scale trend often has a large amplitude that will dominate the repeat abundance in the Fourier transform. To remove this large-scale trend, we first fit a cubic spline curve to the original 1D DNA walk (Figure 1B). We then smooth this trend by subtracting this curve from the original 1D DNA walk (Figure 1C). This is done using the smooth.spline function in the R stats package (smooth.spline function - RDocumentation). This allows the Fourier transform to focus on the local repeats and not the large-scale trends/waves in the DNA walks.

### Fourier transform

A Fast Fourier Transform (FFT) is run on the smoothed 1D DNA walk. The FFT is run on windows of 5000 bp at a time to allow for the detection of a wide range of repeat lengths (from 2bp to 2500bp). We used the fft function in the R stats package (RPubs - Introduction Fast Fourier Transform in R). The FFT outputs complex values. We plot the modulus of these values to produce a strength for each discrete frequency (1/repeat length) (Figure 1D). Large strengths imply a relatively high abundance of the given repeat length in that 5000 bp window. To visualize an entire chromosome’s repeats these 5000bp windows can be joined and relative abundances of each repeat length can be plotted as a heat map (Figure 1E).

These Fourier transforms of 1D AT walks can infer tandem repeats not only in AT but also other nucleotide combinations (e.g. CG, AG, GT, etc.) based on gaps in the AT. There are options within the R package to run the Fourier transform on any pair of base pairs (although AT is the default). Fourier transforms of AT walks were compared to Fourier transforms of CG walks. The same fundamental repeat lengths at the same positions in the genome were detected in both transforms. However, occasionally the abundance of harmonics can vary between the Fourier transforms run on the AT and CG walks. Running 2D wavelet analyses results in complex 2D transformations that are more difficult to interpret and visualize. Here we use 1D walks since we can show that they are able to detect the same fundamental repeat lengths and are easier to understand and visualize. Rare cases where repeats occur only in CG is a situation where running the CG Fourier transform would allow for the specific CG repeat lengths to be determined. Note that these regions will still appear in the AT Fourier transform as dark vertical bars (with no obvious repeat lengths).

### Genomic Shannon diversity

To explore repeat diversity across the chromosome, we introduce a genomic form of the Shannon diversity index (Shannon, 1948; Good, 1953) often used in ecology, where each repeat length is the equivalent of a species, and each window of the genome is the equivalent of a geographic location. The abundance of each repeat output from the Fourier transform is the equivalent of the number of individuals of a given species in a location. Shannon diversity was calculated in R on the Fourier transform output using the diversity function in the vegan package (Oksanen, 2022; diversity function - RDocumentation). The Fourier transform abundance matrix was normalized to values between zero and one by dividing each value in the matrix by the total summed abundance for each repeat length. The Shannon diversity index can be run on the default 5 kbp Fourier windows to find small regions containing densely clustered repeats or on larger (up to 5 Mbp windows) to detect large scale changes in repeat diversity.

### Histograms showing the number of repeat lengths that reach minimum abundance in each window of the genome

As an alternative way to find regions of low repeat diversity (often associated with centromeres), we determine where in the chromosome the most repeat lengths have minimum abundance. Calculating these abundance sum minima involves first finding where in the chromosome the summed abundance of each repeat length is minimized (to within a 2Mbp bin) (findpeaks function - RDocumentation). Next the number of repeat lengths that are minimized in each 2 Mbp bin are counted. The counts of repeat lengths that have a minimum abundance at that location are plotted in 2 Mbp bins (hist function - RDocumentation).

## Results and discussion

RepeatOBserver can be run on any chromosome-scale assembly to visualize tandem repeats, identify centromeres and other regions of potential interest (e.g. neocentromeres, historic centromeres and inversions). Entire chromosomes are represented as 1D and 2D DNA walks, which can be useful for visualizing the dominant base pairs in each region of the genome (Figure 2 A-C). Fourier transforms of AT DNA walks are then used to visualize the repeat spectra in 5, 000 bp windows as a heat map of the abundance of repeats of different size across each chromosome (Figure 2 D and E). Large regions of tandem repeats appear clearly as bright horizontal lines, against a dark background, (e.g. Figure 2D 15-17 Mbp). These bars describe a particular repeat spanning many 5, 000 bp Fourier transform windows along the chromosome. The fundamental frequency detected by the transform is the true repeat length and the lower bars on the plot represent its harmonics. These plots make it easy to visualize repeat patterns across the chromosome.

**Figure 2:**
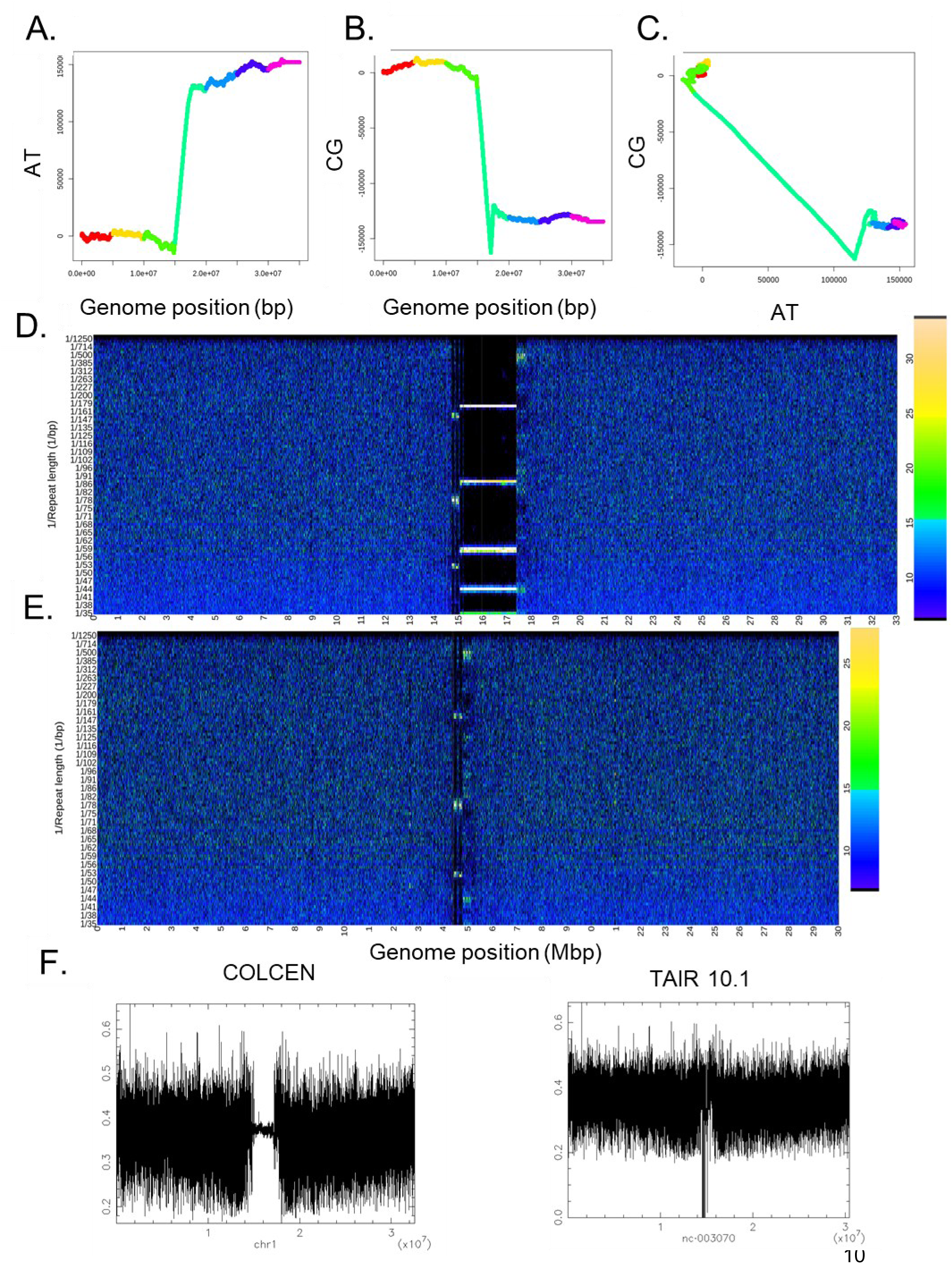
*Arabidopsis thaliana* genome (COLCEN) chromosome 1 (Naish *et al*., 2021) (A) Full chromosome 1D AT DNA walk. (B) Full chromosome 1D CG DNA walk. (C) Full chromosome 2D DNA walks. DNA walks are coloured based on the rainbow; starting at red going to violet and changing colour every 5 Mbp to help visualize how the plots relate to each other. (D) Full chromosome heat map showing the relative abundance of repeats of different lengths in 5 Kbp intervals. (E) Heat map of the older *A. thaliana* TAIR10.1 assembly chromosome 1 (Berardini *et al*., 2015). (F) EMBOSS plots of CG isochores across chromosome 1 in the new COLCEN and older TAIR10.1 assemblies. Comparison between the older *Arabidopsis thaliana* TAIR10.1 assembly (Berardini *et al*., 2015) to the newer COLCEN genome assembly (Naish *et al*., 2021) shows the improvements in assembly quality for repeat sequences in recent years.

Here we show how RepeatOBserver can be used to: (1) visualize repeat patterns across the chromosome (2) identify centromeres, (3) explore variation in repeat patterns across species, and (4) investigate repeat patterns at neocentromeres, areas with gene copy number variation, and inversion boundaries, using a wide range of published plant and animal genomes (Supplementary Table S1).

### Centromere position prediction

There are no bioinformatics programs, to our knowledge, that can accurately and precisely predict natural centromere positions purely from the underlying genomic sequence. While centromeres are defined at least partly epigenetically, they often occur in very repetitive regions in the genome. These regions are usually characterized by the presence of a single dominant tandem repeat found at very high frequency, or the high abundance of a single transposable element. As a result, the centromere position in a chromosome can be consistently identified as the region of the chromosome with the lowest level of repeat diversity. Consistent with this, using RepeatOBserver, we observe a high diversity of Fourier repeat lengths in the non-repetitive, complex regions found in most of the genome, while centromeres show the least diversity in Fourier repeat lengths across a chromosome even when compared to other repetitive parts of the genome. In EMBOSS CG isochore plots, this lack of diversity can often be seen as a steady and often shifted CG isochore signal. In RepeatOBserver heatmaps (e.g. Figure 3A, 5-7Mbp), this lack of diversity often appears as a dark vertical bar with a few bright horizontal frequencies representing the centromeric repeat lengths. In some cases, low repeat diversity in centromeres occurs without any obvious dominant tandem repeat and may be related to an abundance of a transposable element or other semi-repeating sequences. Although centromeres are both genetically and epigenetically determined, we find that natural centromere positions in most of the chromosomes that we explored could be determined based on the repetitiveness of the underlying genetic sequence without the need for epigenetic information.

**Figure 3:**
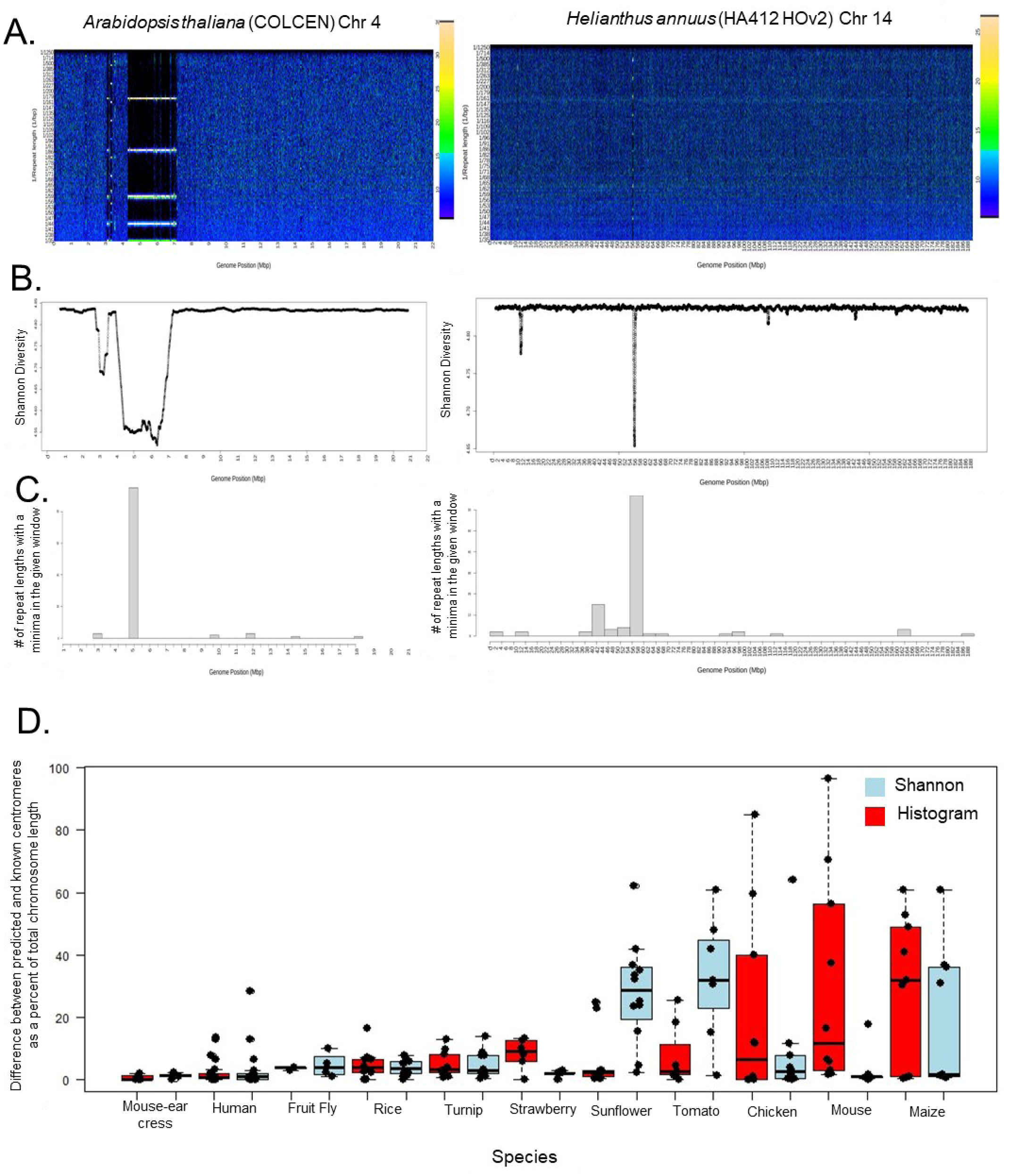
*Arabidopsis thaliana* genome (COLCEN) chromosome 4 (Naish *et al*., 2021) (left) and *Helianthus annuus* (HA412 HOv2) chromosome 14 (Huang *et al*., 2023) (right). (A) Heat map showing repeat lengths and their abundances across the chromosome. (B) Shannon diversity of repeat lengths in each 5 kbp window plotted with a rolling mean over 100 windows. Shannon diversity minimizes at the known centromere position. (C) Histograms show predictions of centromere positions based on where the most repeat lengths minimize their abundance sum over a 2Mbp window. (D) Boxplots showing the error as a percent of the total chromosome length in predicting the centromeres using both Shannon diversity and the histograms across several species. Chromosomes are the black dots, Shannon diversity is plotted in red and the histograms are plotted in light blue. Species included are: Mouse-ear cress - *A. thaliana* (COLCEN assembly) (Naish et al., 2021), Human - *Homo sapiens,* Fruit fly - *Drosophila pseudoobscura lowei* (Bracewell *et al*., 2019), Rice - *Oryza sativa* (Japonica Group) (Nawaz *et al*., 2014), Turnip - *Brassica rapa* (Zhang *et al*., 2023), Strawberry - *Fragaria vesca* (Zhou *et al*., 2023), Sunflower - *H. annuus* (HA412) (Huang et al., 2022), Tomato - *Solanum lycopersicum* (Su *et al*., 2021), Chicken - *Gallus gallus domesticus* (Piégu *et al*., 2018), Mouse - *Mus musculus* (Hughes *et al*., 2007), and Maize - *Zea mays* (Jiao *et al*., 2017).

Given this pattern, we can predict the centromere position in each chromosome by running two analyses (Supplementary Table S2, Figure 3B and C). (1) We calculate a genomic form of the Shannon diversity index (Shannon, 1948); this index is typically used in ecology to look at species diversity in different geographic locations (Good, 1953), but we used it here to explore diversity of repeat lengths across the genome. Using this method, we observed that the absolute minima in repeat diversity across the chromosomes was usually associated with the centromere position (Figure 3B). (2) We also calculate the abundance sum minima across 2 Mbp windows for each repeat length. The number of repeat lengths that have minima in each window can be counted and plotted as a histogram; in what follows, we refer to this as the “histogram method”. The genome window that has the most repeat lengths at minimum abundance is the centromere in most chromosomes. The predicted centromere location is the start position of the 2 Mbp window where this count of minima is largest (Figure 3C). Since telomeres often had the very lowest diversity on this scale, we excluded the first and last 2 Mbp bins from this histogram centromere analysis.

To assess the reliability of these approaches, we compared RepeatOBserver’s predicted centromere locations to the known centromeres of 107 chromosomes across 11 species using both the Shannon diversity and histogram methods (Supplementary Table S3). Centromeres vary tremendously in length between species, which introduces uncertainty when comparing exact locations. The Shannon diversity position is determined by the center of the lowest rolling mean 2.5 Mbp window and the histogram position is returned as the start of the 2 Mbp bin with the most repeat lengths minimized in it. In many cases, we inferred known centromere positions from figures in papers, introducing further uncertainty.

Overall, we found that RepeatOBserver identifies most regional centromeres to within 2-3 Mbp of previously reported positions. This is a close match given that our methods use ∼2Mbp bins, which limits precision. Centromeres in all chromosomes of *A. thaliana*, *Drosophila pseudoobscura lowei*, *Oryza sativa*, *Brassica rapa*, and *Fragaria vesca* were determined within <15% of the total chromosome length by both analyses (histogram and Shannon diversity) using RepeatOBserver (Figure 3D).

The histogram method of centromere identification works even for *Helianthus annuus* and *Solanum lycopersicum* which had centromeres that were not accurately identified by our default Shannon diversity method (Supplementary Table - Centromeres Total). However, for acrocentric centromeres (e.g. *Gallus gallus domesticus*, Mus *musculus* and some *Homo sapiens* chromosomes) only the Shannon diversity method worked. The histogram method was unable to find the centromeres in acrocentric chromosomes since the telomeric regions were excluded. We needed to exclude the telomeric regions from the histograms because nearly all repeat lengths reach an absolute minimum at the telomeres and only a local minimum at the centromeres. Some *Zea mays* centromeres have moved in recent evolutionary time and are known to lack tandem repeats (Talbert and Henikoff, 2020) perhaps explaining why we do not have much success at predicting centromere positions using either method in this species.

In the default Shannon diversity, the majority of known centromeres were correctly identified by calculating the repeat diversity minima in 5 kbp windows and finding the chromosome position where a rolling mean of 500 windows minimized. This method did not need to exclude the telomeres because the telomeres did not minimize their diversity the same way the centromeres did, allowing for telocentric centromeres to be more easily identified. However, for some specific species (e.g. *H. annuus*, *S. lycopersicum*) very different centromere locations are predicted using Shannon diversity depending on the scale of the repeat diversity explored. The true centromere positions in these species show up as local minima using 5 Mbp window scales with telomeres at this scale appearing as the absolute minimum. On the default 5 kbp scale, which successfully predicts centromeres in most species, the absolute and most local minima correspond to other repeating features such as inversion boundaries, subtelomeres and other tandem repeats.

### Comparing RepeatOBserver output to known tandem repeat patterns

To confirm that Fourier transforms of an AT DNA walks can accurately describe genome-wide repeats patterns, we use it on genome assemblies containing well-studied patterns of tandem repeats. When applied to the *Homo sapiens* Y chromosome, RepeatOBserver can easily identify the boundaries of the centromeric, euchromatin and heterochromatin regions (Rhie *et al*., 2023) (Figure 4). In *D. pseudoobscura lowei* chromosomes (Bracewell *et al*., 2019) we find a clear signal at 20-21 bp, matching the known centromeric repeat (Figure 5A). Similarly, we can detect the ∼52 bp repeat known to be associated with centromeric regions in the *S. lycopersicum* genome assembly (Jo et al., 2009) (Figure 5B). In *A. thaliana*, RepeatOBserver plots permit clear identification of the 178 bp repeats (CEN178) that represent the sites where centromere-specific histones (CENTROMERIC HISTONE3, CENH3) are loaded (Wlodzimierz *et al*. 2023). We also detect the known ∼150bp and ∼500bp repeats that precede and follow CEN178 repeats, respectively (Wlodzimierz, Hong, et al., 2023). Interestingly, comparisons between an older *A. thaliana* genome assembly (TAIR 10.01; Berardini *et al*., 2015) and a more recent assembly (Col-CEN; Naish *et al*., 2021) highlight the improvements in the assembly of structural variants and repeated regions that can be obtained with long read technologies.

**Figure 4:**
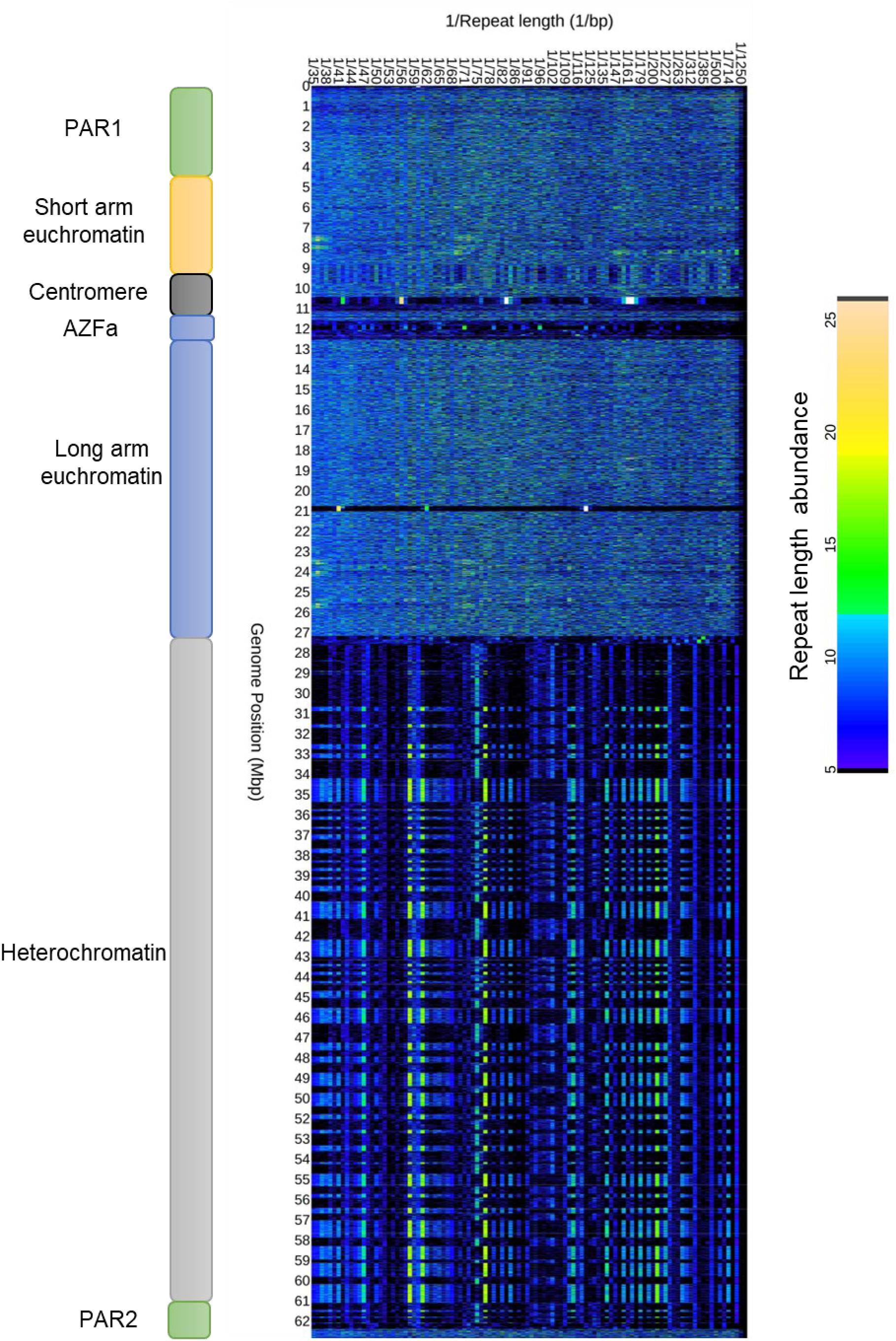
*Homo sapiens* (human) Y chromosome heat map (rotated) showing the relative abundance of each repeat length at each chromosome position. Transitions in repeat structure match known positions in the chromosome for pseudo autosomal region (PAR), euchromatin, heterochromatin and the centromere.

**Figure 5:**
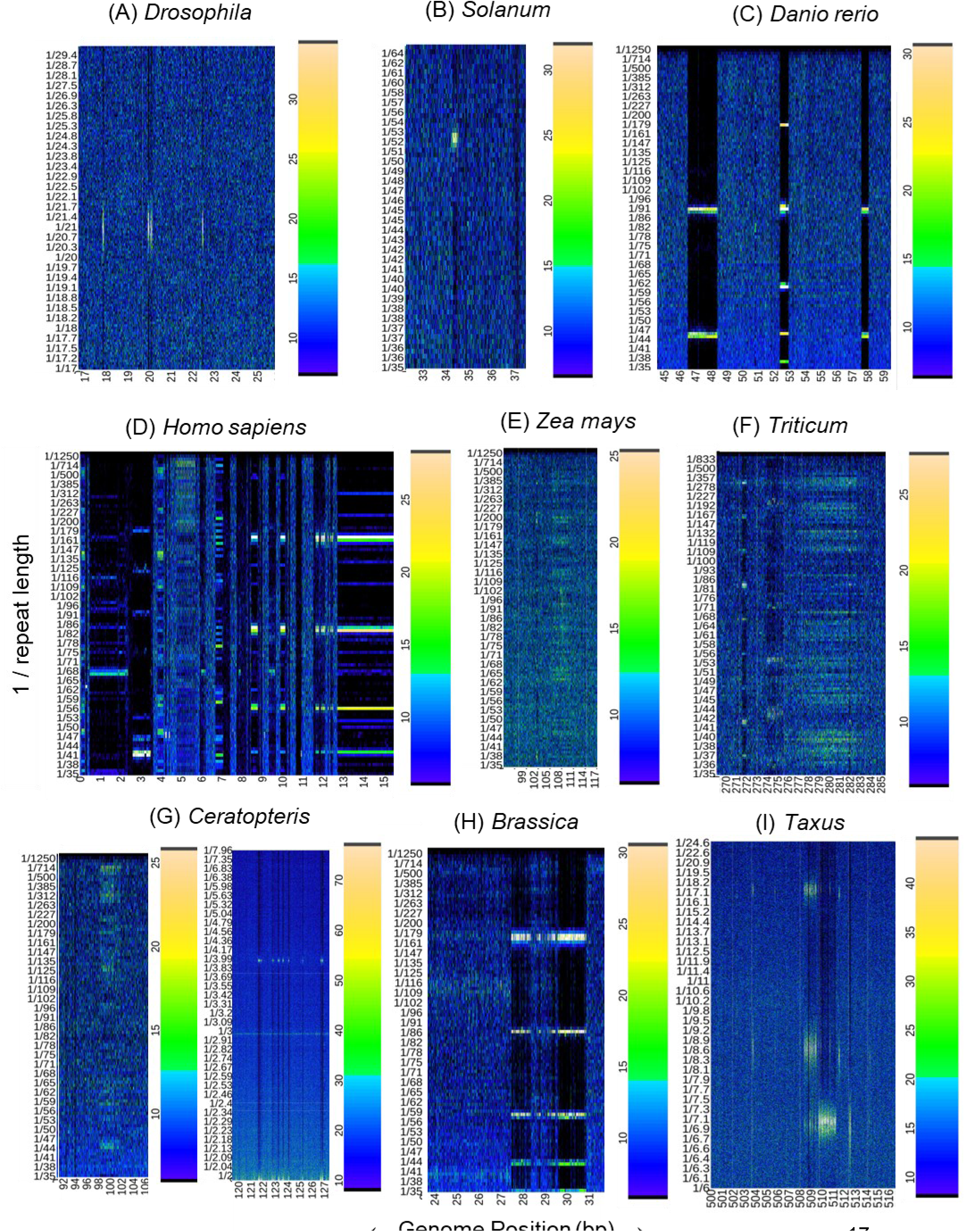
Centromeric repeats shown as heat maps of the relative abundance of each repeat length for (A) *Drosophila pseudoobscura lowei* (Fruit fly) ChrAD, (B) *Solanum pimpinellifolium* (tomato) Chr1, (C) *Danio rerio* (zebrafish) Chr3, (D) *Homo sapiens* (human) Chr22, (E) *Zea mays* (maize) Chr 4, (F) *Triticum monococcum* (einkorn wheat) Chr 4A, (G) *Ceratopteris richardii* (fern) Chr 2, (H) *Brassica rapa* (turnip) Chr 5 and (I) *Taxus wallichiana* var*. yunnanensis* (yew) Chr 3.

EMBOSS plots (Rice *et al*., 2000) of CG isochores provide a simple means for detecting the centromeres in many species with AT rich centromeric sequences. However, in some plant and animal genome assemblies it is less clear if there is always an isochore signal at the centromere (e.g. *H. annuus*). We were not able to determine whether this lack of signal is just related to the quality of the reference genome (e.g., newer vs. older *A. thaliana* chromosomes in Figure 3F) or to the actual composition of repeats throughout that genome. We hypothesize that it may be related to the type of centromere, with satellite-rich, AT-dense old centromeres appearing clearly in CG isochore plots and recently originated centromeres characterized by retrotransposons not appearing at all (e.g. in *Z. mays* or *H. annuus*).

### Variation in tandem repeat patterns

By plotting the Fourier transforms in a wide variety of species, we can begin to visualize the diversity in repeat patterns that exist across the chromosome and in the centromere specifically (Figure 5). Both *D. pseudoobscura lowei* and *S. pimpinellifolium* are examples of species that have single regional centromeres on most chromosomes. They are consistently represented by a known repeat of length 21 bp in *D. pseudoobscura lowei* (Bracewell *et al*., 2019) and 52 bp in *S. pimpinellifolium* (Jo *et al*., 2009) (Figure 5 A and B).

Many other species’ centromeres contain tandem repeat patterns of varying lengths (Figure 5C). These different tandem repeats will appear in the Fourier spectra as bright horizontal bars at different positions (the top bar is the fundamental frequency or true repeat length and the bars under it show higher-frequency harmonics that describe the complexity of the specific repeat). Each fundamental frequency and its harmonics represent a particular kind of repeat pattern. The same horizontal bar pattern in two locations implies the same repeat at two different positions in the genome (e.g. a fundamental repeat that is 90 bp long with a harmonic at 45 bp can be seen at both 47 Mbp and 57 Mbp in *Danio rerio* (zebrafish) chromosome 3 in Figure 5C). In *D. rerio*, it is evident that these matching repeats are separated by a distinct repeat with a different fundamental length (180 bp long) and harmonics at 1/2 (90 bp), 1/3 (60 bp), 1/4 (45 bp) and 1/5 (36 bp). If there is a lower fundamental frequency (large repeat) that is too long for the 5000 bp window that we used, then this pattern would be the fraction of the true fundamental repeat length. The true fundamental could be calculated (or determined using a larger Fourier window, e.g. run_long_repeats function using a 20 kbp window option in R package). Metacentric (centered on the chromosome) centromeres are the easiest to detect even if they have more than one fundamental repeat frequency.

### Subtelomeric repeats and acrocentric centromeres

In many mammal and bird genomes, chromosomes are acrocentric. In *H. sapiens* chromosomes 13, 14, 15, 21, 22, X, and Y, an extended set of complex repeats marks the subtelomeric region and centromere. Acrocentric centromeres can be difficult to tell apart from these subtelomeric repeats (e.g. *H. sapiens* Chromosome 22, Figure 5D). In some chromosomes these sub-telomeric repeats are even less diverse (using Shannon diversity) than the centromeric repeats making it difficult to consistently differentiate between centromeres and subtelomeric repeats.

### Retrotransposons

Clusters of retrotransposons can be detected using RepeatOBserver and appear as vertical ‘blurs’ in the Fourier transform. Some centromeres do not appear to be composed of one or many clear tandem repeats and their harmonics; instead, they are made up of clusters of retrotransposons. *Zea mays* (Dawe *et al*., 2023) and *Triticum monococcum* (bread wheat and einkorn wheat (Ahmed *et al*., 2023)) are examples of such centromeres (Figure 5E and F). The blur in the Fourier transform implies that the repetitive sequences vary considerably in length. In *T. monococcum*, it was found that the centromeric regions contained many retrotransposons (Ahmed *et al*., 2023) suggesting that these blurs in the Fourier spectra are picking up the repeated insertions of retrotransposons that are near each other but not perfectly in tandem; hence the variation in lengths. Past studies have suggested that this lack of clear tandem repeats is related to neocentromere formation since it is often observed in domesticated species with recently moved centromeres (Talbert and Henikoff, 2020). A similar pattern appears in some *Ceratopteris richardii* (fern) chromosomes suggesting they may also have centromeres composed of retrotransposons.

### Periodicity

DNA periodicity is also evident when looking at the Fourier spectra. In all chromosomes, there is a well-known 3 bp periodicity in exons that is evident in the Fourier spectra of all chromosomes (Eskesen *et al*., 2004; Nagai *et al*., 2001). In *C. richardii* chromosomes, there is also a 4 bp repeat that appears at semi-regular intervals throughout all the chromosomes (Figure 5G). Future studies should explore if these repeats mark any important boundaries in the genome. Other periodicities with slight variations in repeat lengths appear as blurs in various genomes and can be visualized using this technique (e.g. ∼19 bp in *Anolis sagrei*, ∼47 bp in *Z. mays*, and ∼60-100 bp and 165 bp in *H. sapiens* (Figure 5D)). In cases where this regular periodicity is blurred it is possibly caused by a high abundance of retrotransposable elements of that approximate length in that region of the genome (e.g. *Brassica rapa* centromere Figure 5H). In *Taxus wallichiana* (yew, Figure 5I) we show a predicted centromere location that has somewhat blurred repeats likely due to the overall imperfections (mutations, insertions, and deletions) in the repeats.

### Neocentromeres, variable number tandem repeats (VNTRs), and gene copy number variation

RepeatOBserver can identify a wide range of perfect and imperfect repeating patterns in DNA sequences throughout the chromosome. While the dominant and often most clear tandem repeats in a chromosome are found at the centromere, it is possible to visualize other repetitive sections of the genome including subtelomeric repeats, neocentromeres, variable number tandem repeats, variation in short tandem repeats and regions containing gene copy number variation. To explore the visualization of these non-centromeric repeats, we chose to look at *H. sapiens* chromosome 8 (Figure 6) which contains a variable number tandem repeat (VNTR) that has been labeled as a neocentromere (85-87 Mbp) as well as three regions (7-7.6 Mbp, 11.5-12.2 Mbp, and 12.2-12.8 Mbp) that contain copy number variation for beta-defensin (a gene involved in flu immune response) (Logsdon *et al*., 2021). Repetitive sequences at all these locations were visible in the Fourier spectra of this chromosome and appeared as weaker bands than the ‘pure’ tandem repeats seen in many centromeres but not as blurred as retrotransposon clusters.

**Figure 6:**
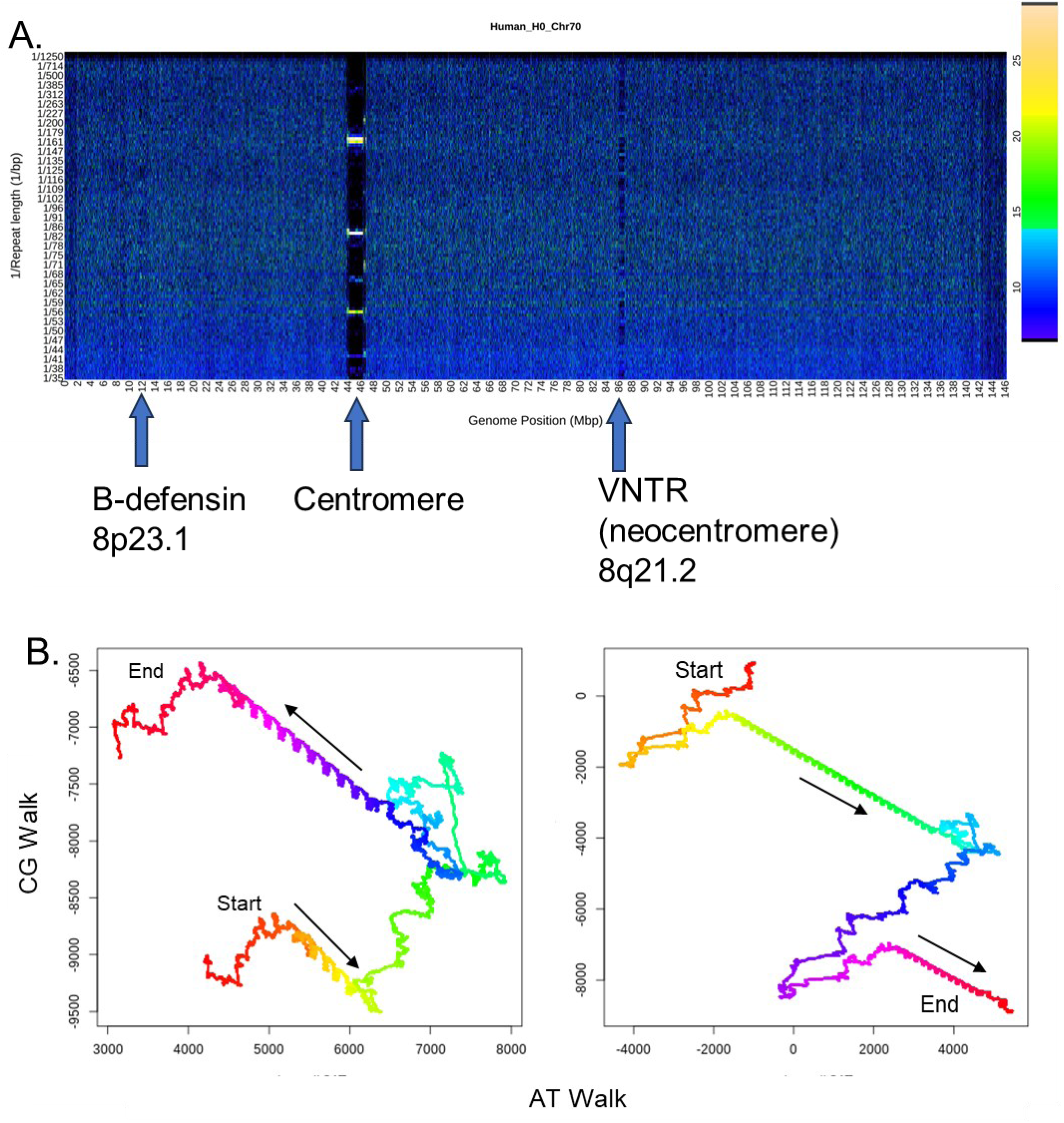
(A) *Homo sapiens* (human) chromosome 8 heat map showing the relative abundance of each repeat length at each chromosome position. The centromere appears as the dominant tandem repeat at ∼45 Mbp. A variable number tandem repeat that has been observed to act as a neocentromere appears at 86 Mbp. Gene copy variation of beta-defensin appears from 7 Mbp to 13 Mbp. (B) This region (7-13 Mbp) contains an inversion that we can visualize in the DNA walks of these sequences. Tandem repeats start at the bottom right and move to the top left in the region from 7-7.6 Mbp and are reversed in the following region (11.5 Mbp) starting in the top left and moving to the bottom right. DNA walks are coloured based on the rainbow; starting at red going to violet and changing colour to help visualize directionality.

### Inversions bounded by tandem repeats

In *Homo sapiens* chromosome 8 described above, the gene copy variation regions are known to be split by an inversion from 7.5 Mbp-11.6 Mbp (Logsdon *et al*., 2021). This inversion can be visualized in the DNA walks of these sequences (Figure 6B and C) since the tandem repeats in these walks are inverted relative to each other on either side of the inversion.

Similarly, in *Helianthus annuus* there is a known inversion on chromosome 5 that differs between two cultivars (HA412 and HA89) for which reference sequences are available (Huang *et al*., 2022). The repeat spectra in HA89 shows the same repeat on both sides of the inversion while in HA412 the repeats are not split by the inversion. When these split repeats were plotted in a 2D DNA walk it was apparent that the first set of repeats in HA89 were in the same direction as the repeats in HA412. However, the second set of repeats in HA89 were clearly inverted relative to those in HA412 and the first set of repeats in HA89 (like the inverted repeats in *H. sapiens* chromosome 8).

### Main features to look for in RepeatOBserver output

When looking at a newly assembled chromosome using RepeatOBserver, it is helpful to initially confirm that the 6-7 bp telomerase repeats are found in at least some chromosome ends. If they are found elsewhere in the chromosome, the possibility of a mis-assembly should be considered. The centromeres can most likely be found where histograms show the most repeat lengths reach a minimum abundance. In this genome position in the heat map plots there will often be bright clear horizontal bars with dark regions in between. The top bar will usually be the fundamental or true repeat length and the bars below it are the harmonics. If the first harmonic below the fundamental is not half of the fundamental, it can be worth it to run the Fourier transform with a larger window size (e.g. 20 kbp) to check for larger repeats. The fundamental can also be calculated from the harmonics, if needed. Blurred bright bars (e.g. Figure 5F, 8A 21-44Mbp) in the heat maps indicate a potentially retrotransposon-dense region. If there is only one such region in the chromosome, this is usually the centromere, even if another region contains more clear tandem repeats (Figure 5F). Gene copy variation and some neocentromeres can appear as anything from perfect to semi-blurred fundamental and harmonic repeat lengths in the heat maps. Matching but separated tandem repeats can potentially indicate an inverted region (Figure 7B HA89 and Figure 8B *Brassica rapa*). If the split repeats are inverted relative to each other in a DNA walk of these repeats, this further supports a potential region to search for inversion signals (Figures 6B and 7C). Holocentric chromosomes will appear as multiple dark vertical bars with similar repeat structure throughout the chromosome (Kuo *et al*., 2022; Figure 8C and D). Not only can RepeatOBserver work well for exploring reference genome assemblies, but it also provides a simple method to visualize repeat patterns in many haplotypes within a species, allowing users to spot mis-assemblies and quickly determine changes to centromere and other tandem repeat structures.

**Figure 7:**
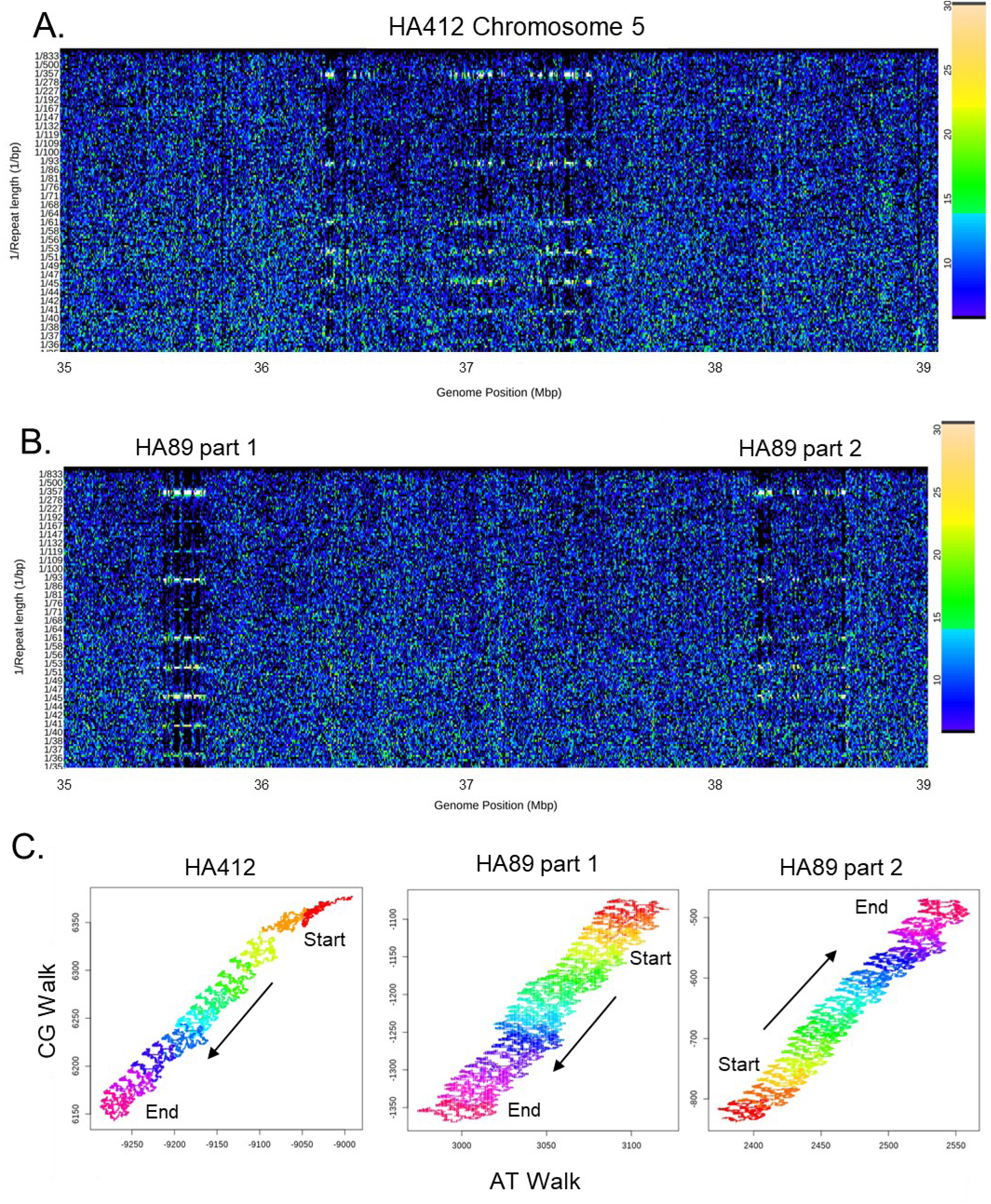
Inversion between sunflower (*Helianthus annuus*) haplotypes HA89 and HA412 on chromosome 5. (A) Chromosome scale Fourier spectra of HA412 showing faint repeats at 27 and 36 Mbp. (B) Zooming in to look more closely at repeats at 36 Mbp in both haplotypes (HA412 and HA89) since there is a known inversion that varies between haplotypes at this location. Repeats are split at the inversion boundaries in HA89. (C) Repeats shown in the DNA walks for HA412, HA89 part1 and then same repeats inverted for HA89 part2. DNA walks are coloured based on the rainbow; starting at red going to violet and changing colour to help visualize directionality.

**Figure 8:**
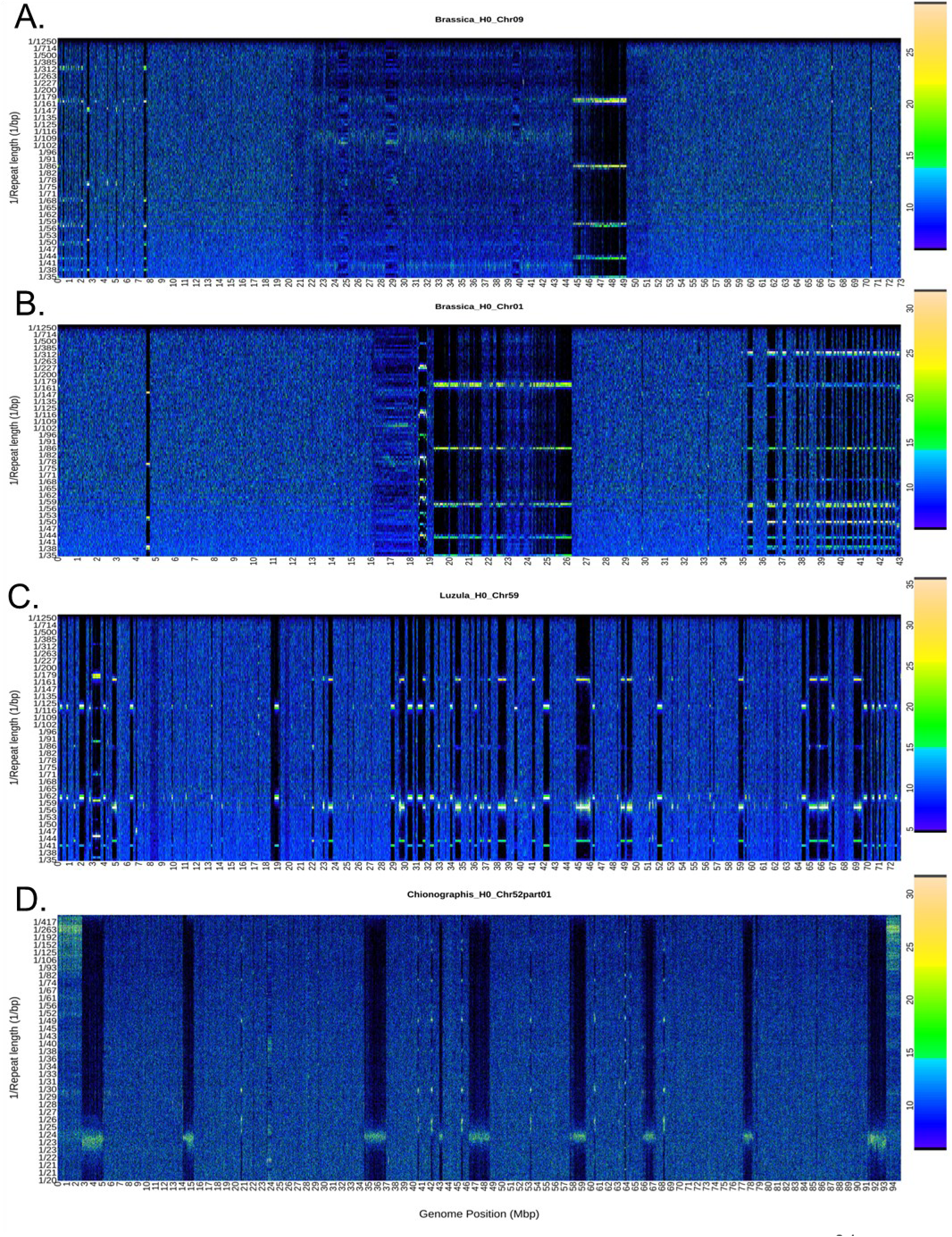
Example repeat spectra showing clear tandem repeats, split repeat regions, different fundamental repeat lengths, potential dense regions of retrotransposons, and potential neocentromeres from (A) *Brassica rapa* (turnip) Chr9 showing clear tandem repeats in centromeres and blurred retrotransposons, which resemble the gene copy number variation and neocentromere in human Chr8 (B) *B. rapa* Chr1 showing split repeats. (C) holocentric chromosomes in *Luzula sylvatica* Chr 4 and (D) *Chionographis japonica* Chr 4 showing the spread out but still clearly visible tandem repeats (Kuo *et al*., 2022).

When searching for the centromere positions, ideally the Shannon diversity and histogram outputs would agree and predict the same centromere location that has also a clear repeat pattern in the heatmap repeat spectra. If the Shannon diversity and histogram outputs do not agree, it is worthwhile to check the repeat spectra heatmaps across multiple chromosomes to determine if the species being explored has a unique centromere structure that appears in multiple chromosomes (e.g., a specific repeat length, due to retrotransposons, acrocentric centromeres, or a recent shift in their centromeres). For species with less common centromeric patterns, we were typically able to successfully predict centromere positions using the heat maps of their Fourier spectra. For example, in wheat we could see a transposable element pattern associated with centromeres appearing once in every chromosome.

Using the default Shannon diversity index to measure repeat diversity across each chromosome, it is possible to not only spot likely centromere positions, but also visualize other changes in sequence repeat diversity that could be caused by small repeat clusters (e.g. trinucleotide repeats) to much larger tandem repeat clusters (occasionally associated with gene copy number variation and inversion boundaries). The amount of imperfection in repeats due to insertions and deletions can be determined for a specific repeat based on the difference in repeat abundance between the fundamental repeat length and the next closest repeat lengths (e.g. for a 50 bp repeat the abundance of 50 bp compared to 51 and 49 bp gives a relative imperfection amount). For base pair changes in the sequence itself, these can be spotted in the similar ‘blur’ of the harmonics but are harder to interpret without finding the specific repeating sequence in the fasta file. Using the Shannon diversity index, we were also able to quickly spot small changes in repeats between varieties of the same species (e.g. a triplet expansion just ∼1000 bp long in one line of *A. thaliana*).

### What evolutionary forces could be driving the lack of repeat diversity at centromeres?

Using RepeatOBserver we can visually observe the different kinds of repeats and repeat patterns characteristic of centromeres that have been reported in many previous studies (e.g. Melters *et al*., 2012; Logsdon *et al*., 2023). Haig (2022) states that centromeres must be ‘hospitable’ environments for repeats. We observe that within a particular chromosome the diversity of repeat types is lowest at the centromere implying that one or a few pure repeats typically play an important role in centromere structure. We can also see, as noted before (Shepelev *et al.,* 2009), that the clearest repeats are often in the middle of the centromeric region and sequences that are less conserved ‘blur’ as you go farther from this central point (Supplementary Figures Fourier heatmaps of all *Homo sapiens* chromosomes). What evolutionary forces are driving the lack of repeat diversity at centromeres while allowing for rapid evolution of centromeres between species and even chromosomes within a species? Many past studies have looked at the evolution of repeats (TEs and tandem repeats) in centromeres (reviewed in Hartley and O’Neill, 2019). It was initially suggested that unequal crossing over generated the varying abundances of repeats around centromeres (Smith, 1976). However, replication slippage and unequal crossing over are unlikely to be the driving factor since recombination is extremely limited around centromeres (Walsh, 1987). Gene conversion (Shi *et al.,* 2010) and interchromosomal and mitotic recombination (Jaco et al., 2008) have been suggested as factors that may be contributing to the rapidly varying copy numbers and similarities seen between some centromeres in the same species. A lack of methylation causing a change in mitotic recombination resulted in a decrease in the repeat copy numbers in mice (Jaco *et al.,* 2008). It is still unclear why centromeric repeats match between some chromosomes and not others within a given species.

There is also uncertainty regarding the origin of centromeric repeats. One of the older hypotheses is the library hypothesis (Salser *et al*., 1976). It suggests that every genome contains a wide variety of repetitive sequences. If one of these sequences ends up in the centromere due to a chromosomal rearrangement or some other shift this repeat will increase in abundance due to the processes described above (Hartley and O’Neill, 2019). Another hypothesis is that centromeric repeats derive from subtelomeric repeats (Villasante *et al*., 2007). Although this may be the case in some species (especially those starting with telocentric centromeres), many studies have reported very fast rates of evolution in centromeres that do not resemble the subtelomeric repeats in that species. The centromere drive hypothesis (Malik, 2009) suggests that some repeat sequences provide a transmission advantage. In response to this, the centromeric proteins (e.g. CEN-A) evolve rapidly to re-balance meiosis. Recently formed centromeres do not always have clear, large tandem repeat arrays suggesting these repeats form later once the new centromere has reached a certain stability in the population (Hartley and O’Neill, 2019). Throughout all the genomes that we explored (including fairly recently formed centromeres in *Drosophila*), we were mostly able to detect the centromere as the genomic location with the lowest diversity of repeats (not necessarily implying a single dominant repeat length but implying a lower genome complexity than found in the rest of the genome). In some chromosomes in both *Triticum* and *Zea mays*, there is a high abundance of retrotransposons but no clear tandem repeats making up the centromere (Ahmed *et al*., 2023). Based on the many types of centromeric repeats and their structure it seems possible that all of these hypotheses contribute to some centromere evolution.

## Conclusions

With the recent advances in long read sequencing technology, genome assemblies more frequently include long stretches of tandem repeats, creating an opportunity to visualize and compare chromosome-scale tandem repeat patterns between species and individuals. Across plants and animals there is enormous variation in tandem repeat structure. RepeatOBserver plots allow for easy visualization of fundamental tandem repeat lengths, repeat blur/clarity, and repeat positions throughout the genome without the user needing to choose any specific input parameters. For chromosomes with a single regional centromere, the centromere appears to consistently be found where all but a few repeats minimize due to the dominance of a particular repeat and its harmonics. Detection is more difficult, but still possible, when centromeres are merged with subtelomeric repeats in acrocentric centromeres or when they are composed of retrotransposons but not tandem repeats. Clear tandem repeat patterns not at the centromere are often associated with subtelomeric repeats, gene copy variation, neocentromeres and potential inversion boundaries. These genomic features appear clearly in the repeat spectra plots highlighting the potential for this tool to visualize and search for these features in newly assembled non-model species. Even for the top model species like *A. thaliana*, current reference assemblies contain large stretches of tandem repeats that were missing from genomes assembled only a few years earlier, making inferences from these patterns much easier now.

RepeatOBserver is a simple and accessible program to detect tandem repeats patterns and centromere positions in a chromosome-scale genome assembly. It requires no input parameters or prior knowledge of the repeats and can even detect imperfect repeats. Although it is ideally run on a server (e.g. ∼20 min on 15 CPU per chromosome), it works on a personal computer given enough time. This program provides a quick way to visualize whole chromosome structures, helping bioinformaticians locate centromeres and other repetitive regions of interest like retrotransposons clusters and gene copy number variation.

## Supplementary materials

Supplementary Table S0: A comparison of bioinformatics tools used for whole chromosome visualization, tandem repeat detection and centromere location prediction.

Table 1 Bioinformatics tools.xlsx

Supplementary Table S1: Links to all the genomes used in this paper. Table S1 Links to genomes explored.xlsx

Supplementary Table S2: Known, histogram and Shannon diversity predicted centromere positions for each species with known centromere locations shown in Figure 3C.

Table S2 Comparing_known_predicted_centromeres.xlsx

Supplementary Table S3: Histogram and Shannon diversity predicted centromere positions for all the species we ran the program on (including several species without known centromere positions).

Table S3 Centromeres_total.xlsx

Supplementary figures: Fourier heatmaps showing repeat lengths from 35-2000 bp in all the human chromosomes to better represent what an entire genome looks like.

Human_Repeats.pdf

## Acknowledgements

This paper is written in memory of Rob Elphinstone. His endless passion and curiosity for mathematics, physics and biology inspired all this work. Thank you also to Debbie Hearn for all her help and support in writing the paper.

