## Supplementary figures and images for "RepeatOBserver: tandem repeat visualization and centromere detection"

### Supplementary figures

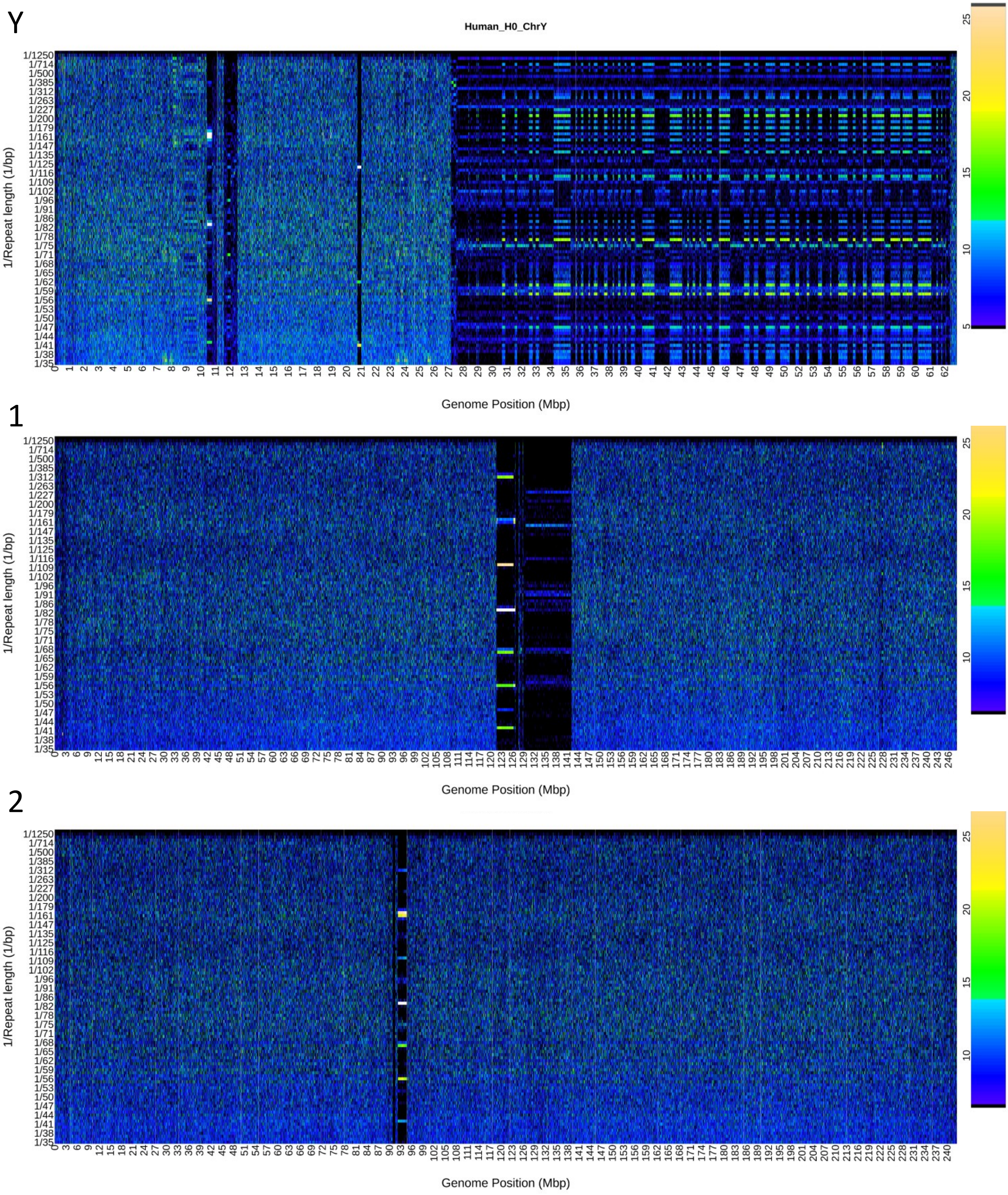

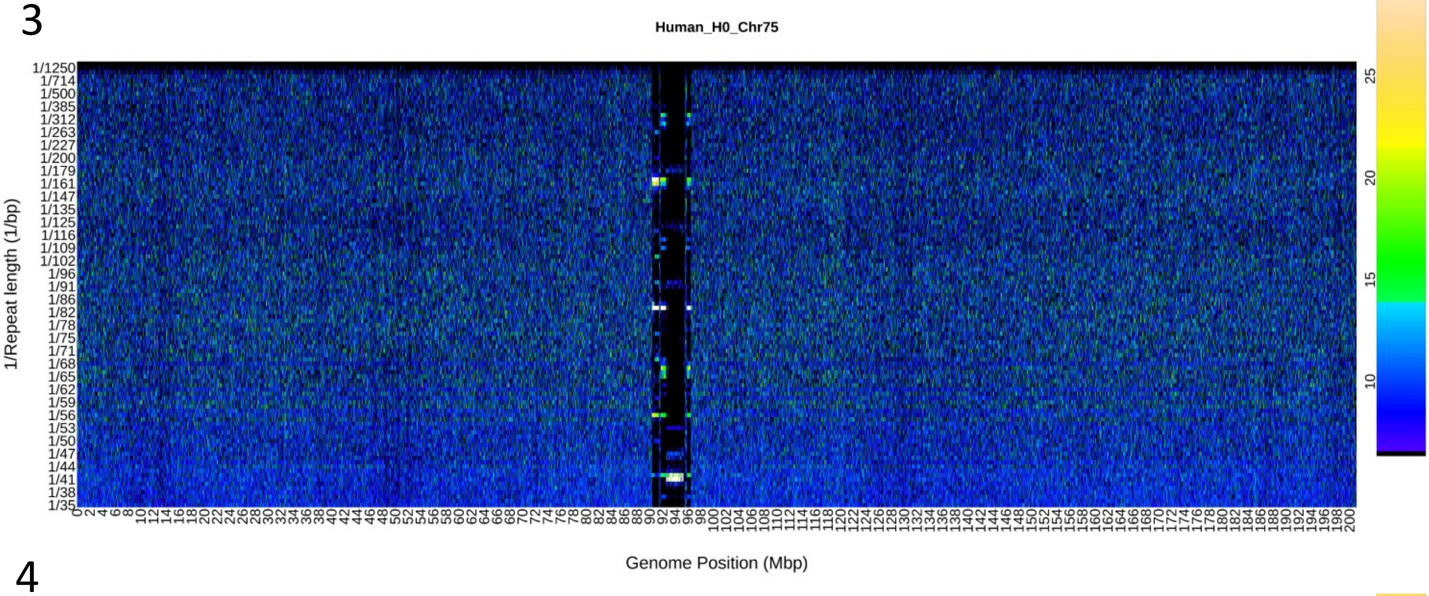

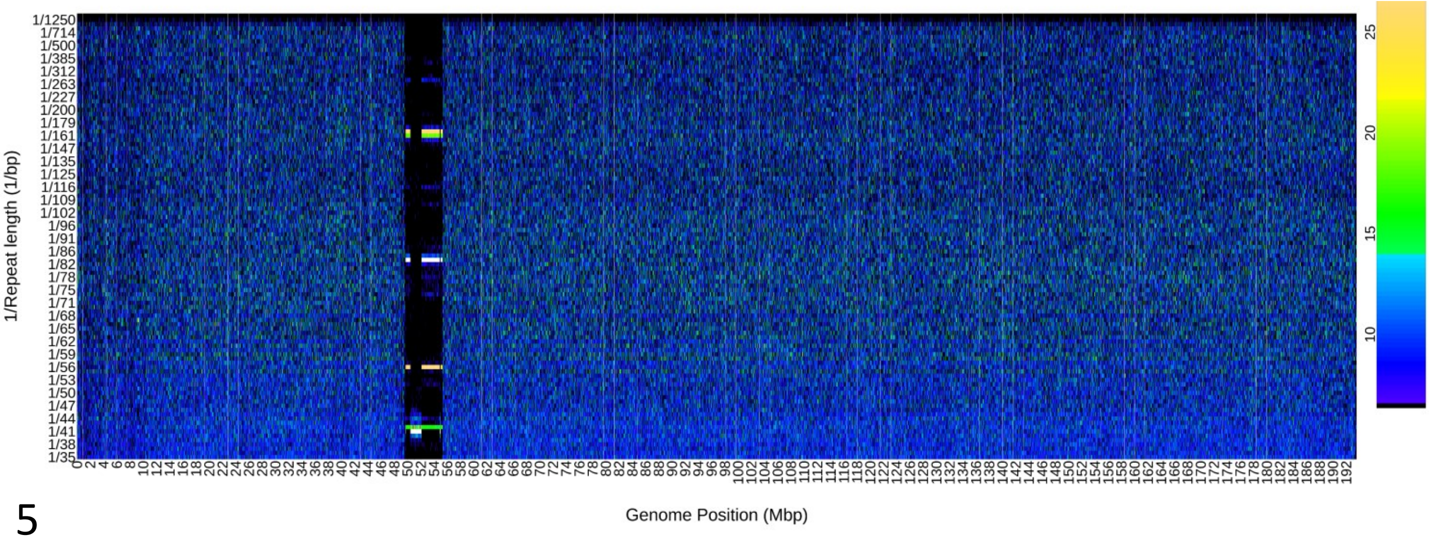

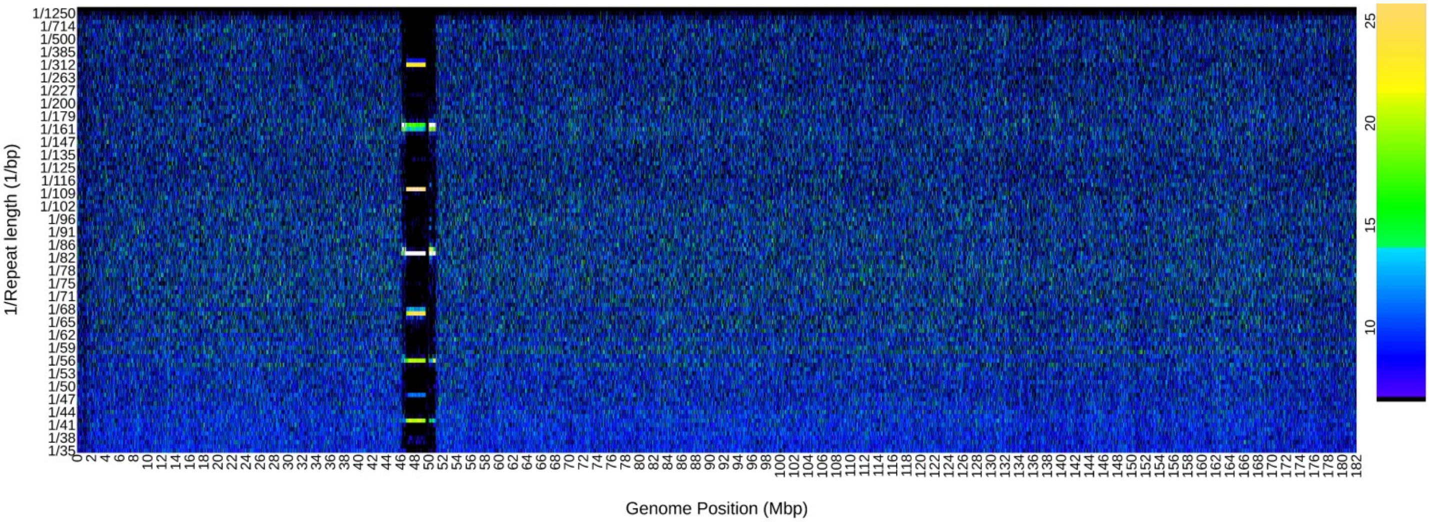

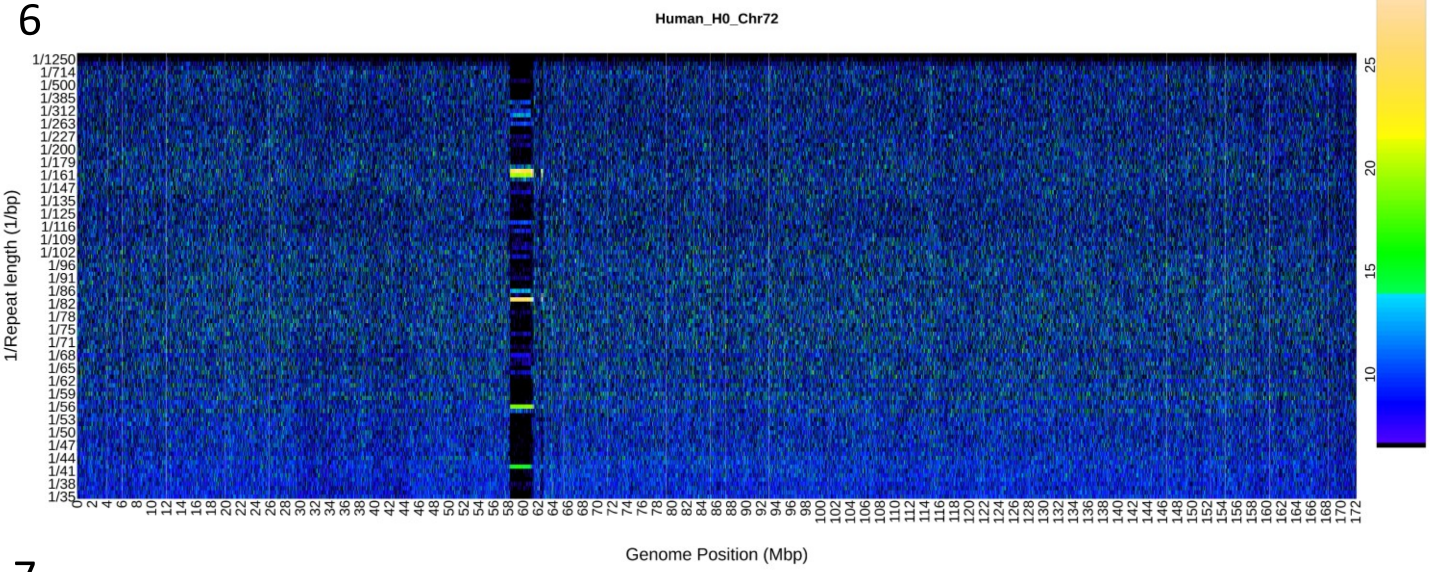

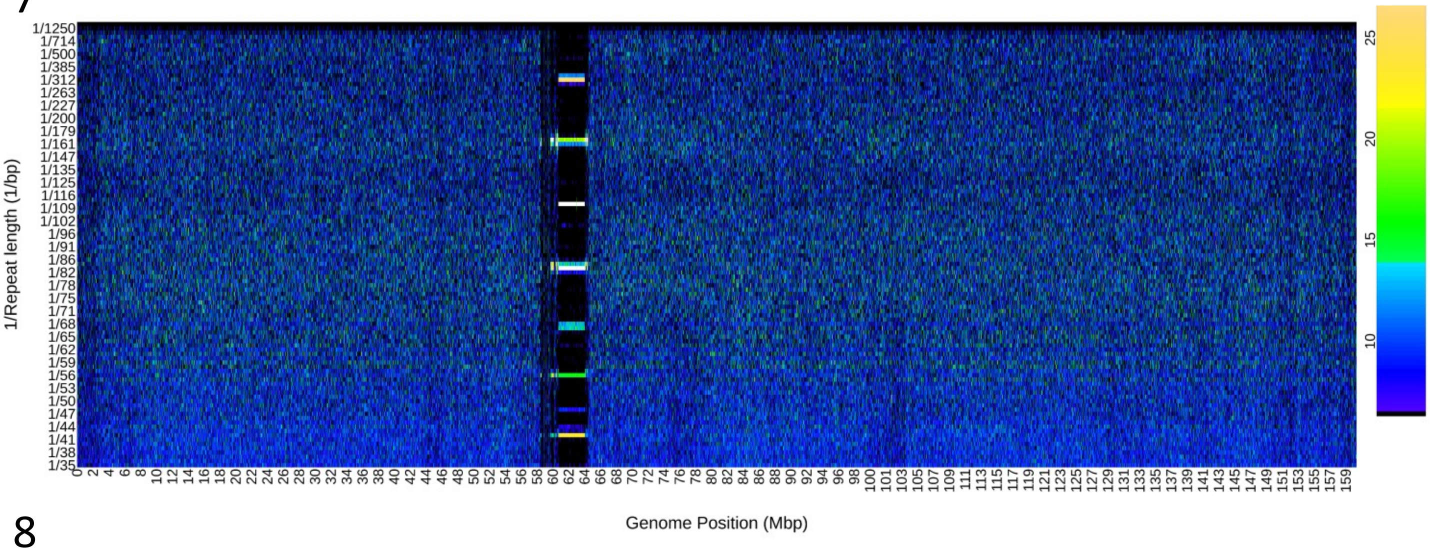

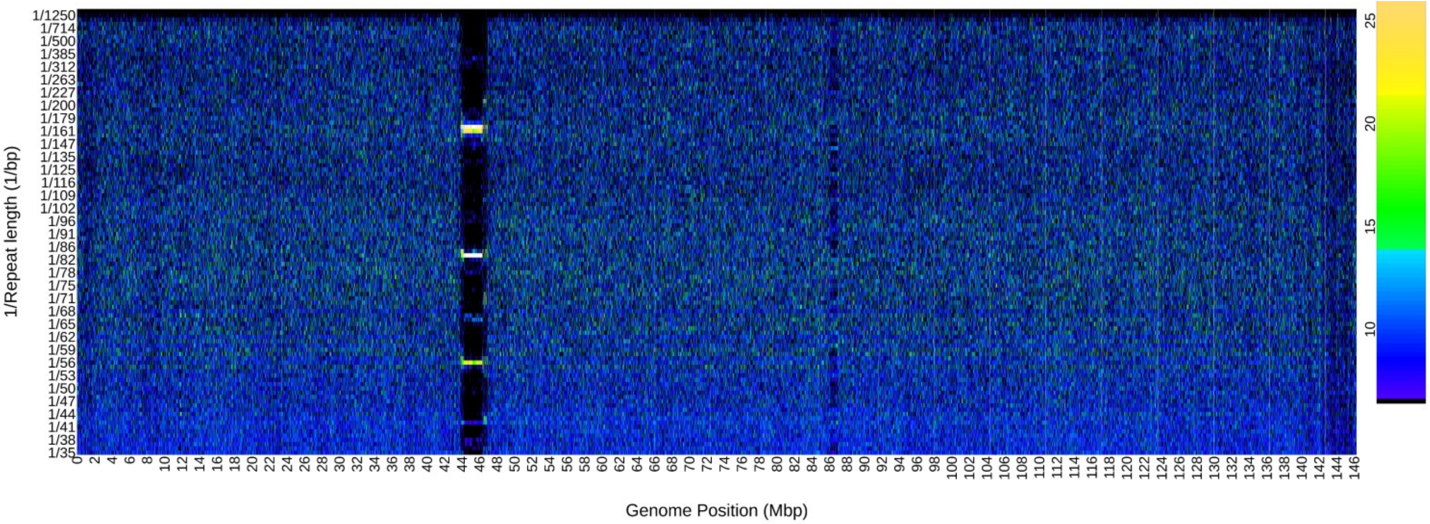

9

Human_H0_Chr69

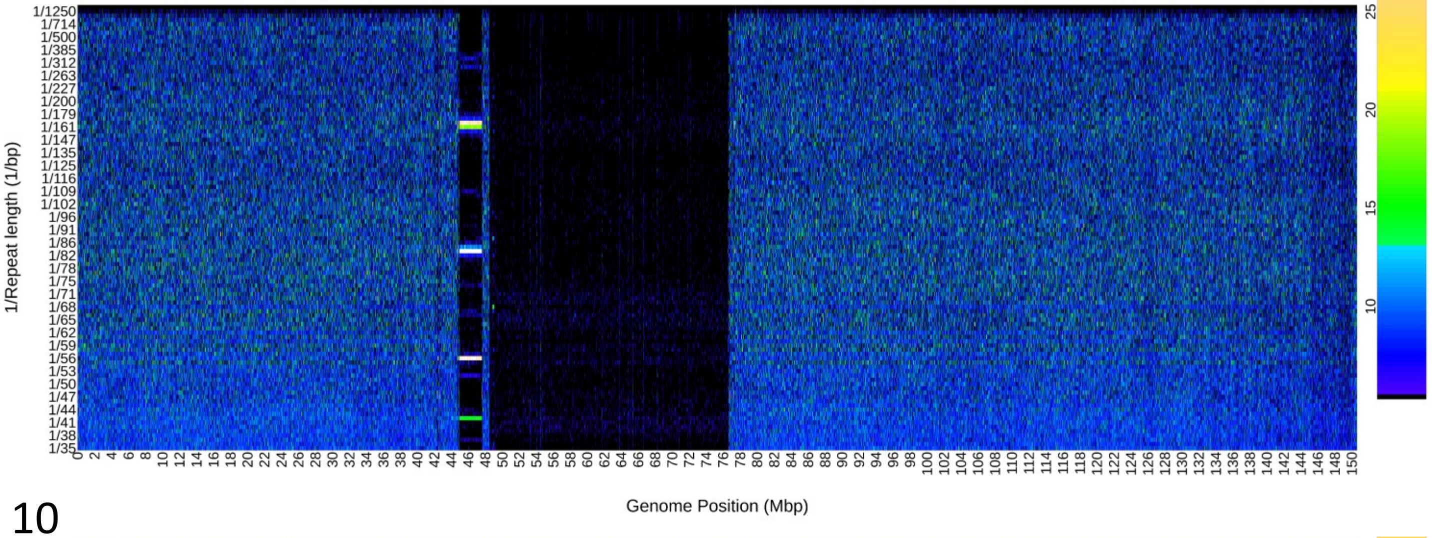


10

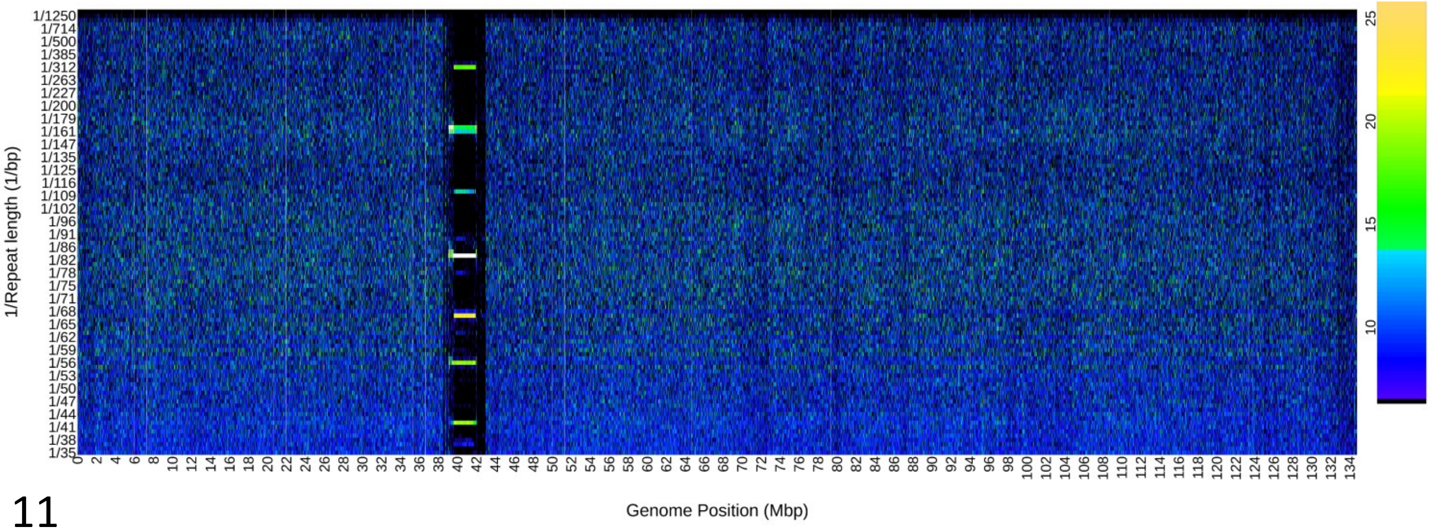


11

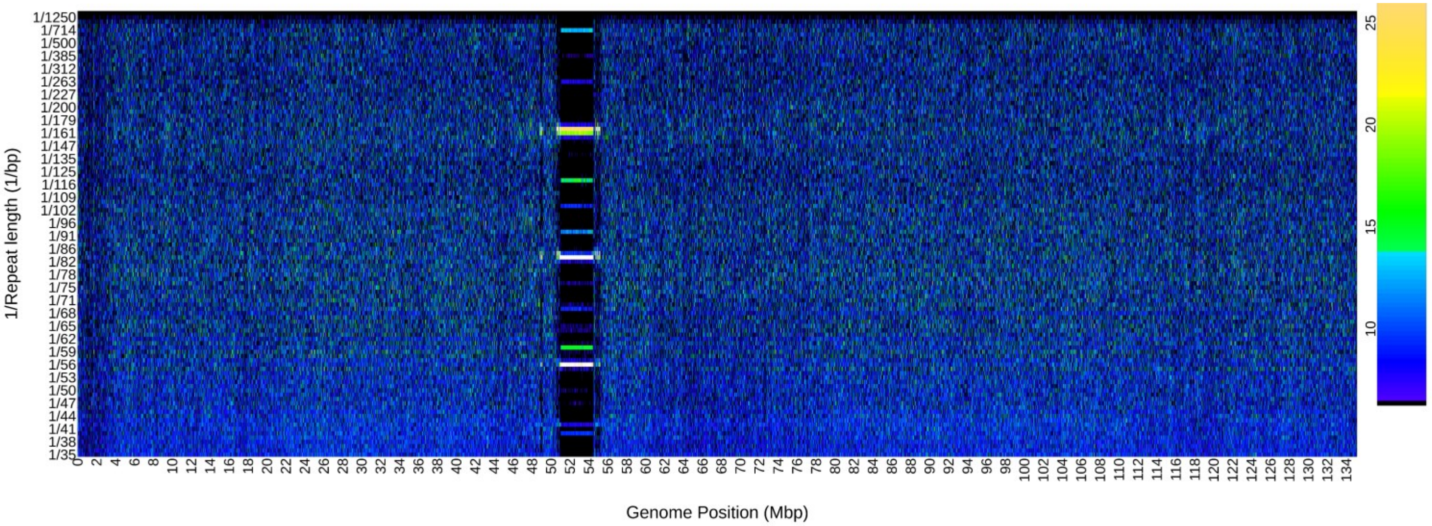

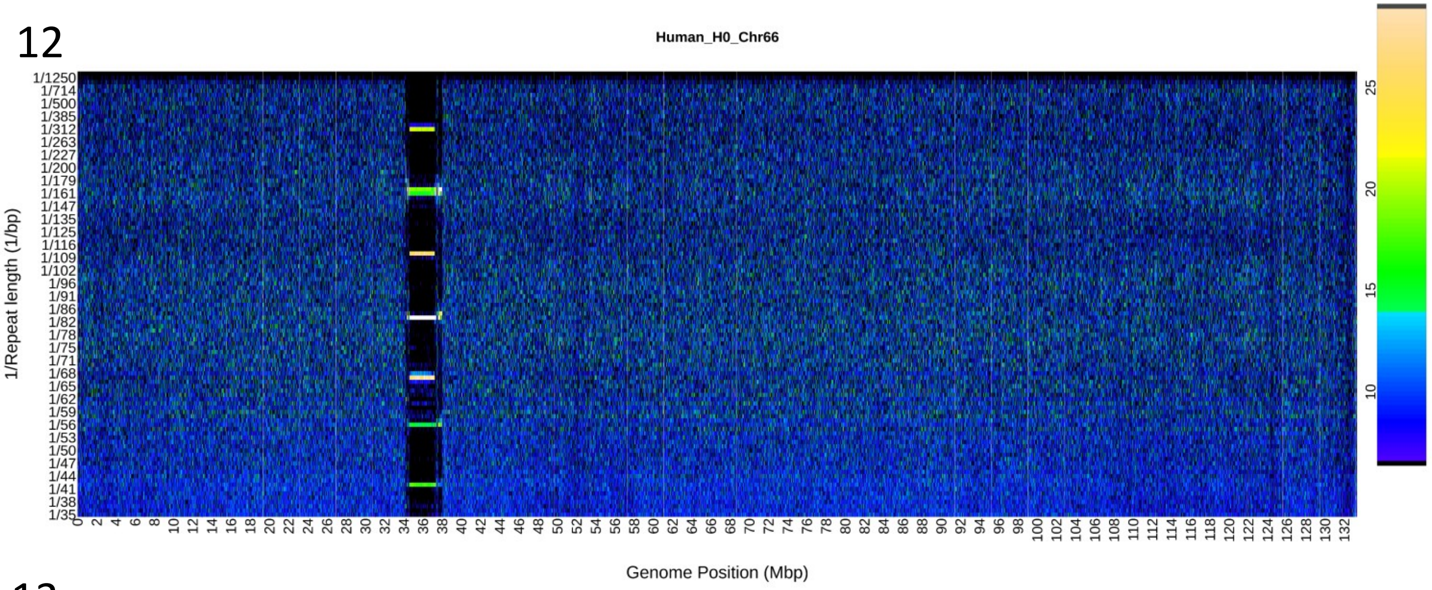

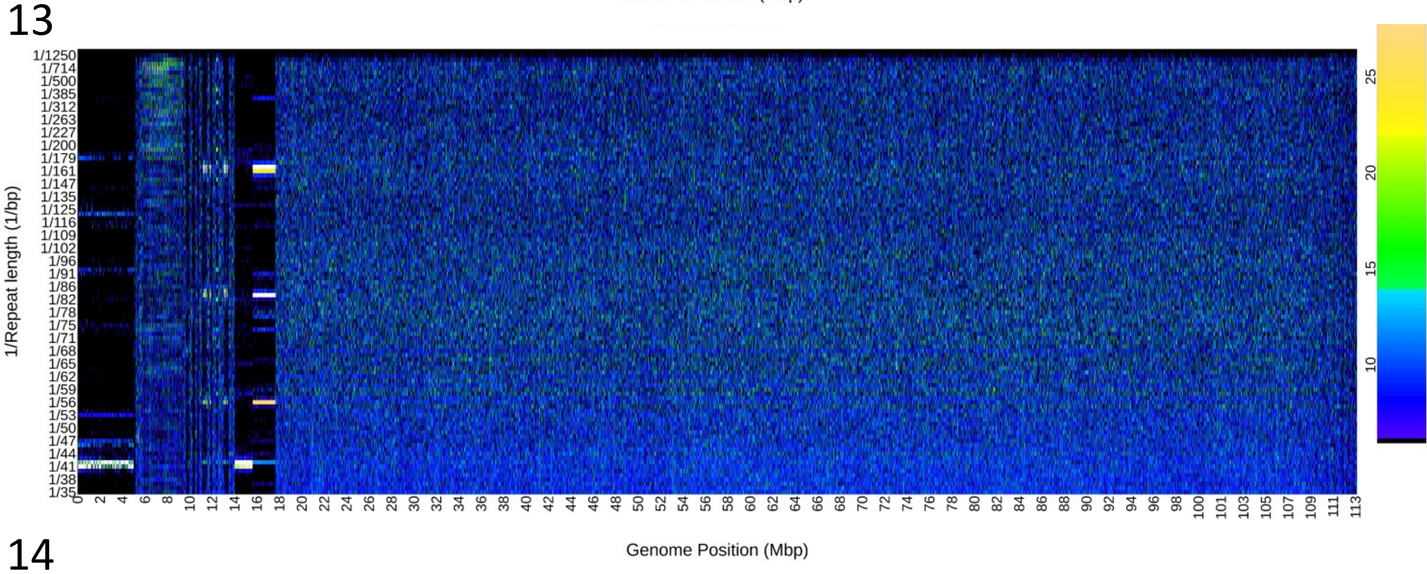

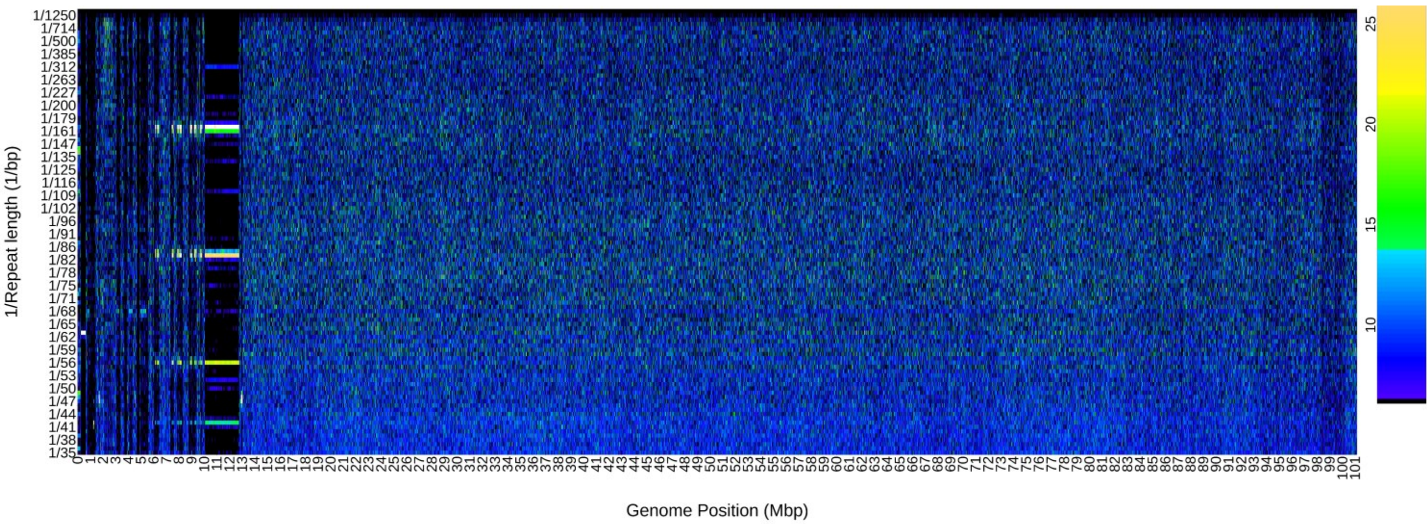

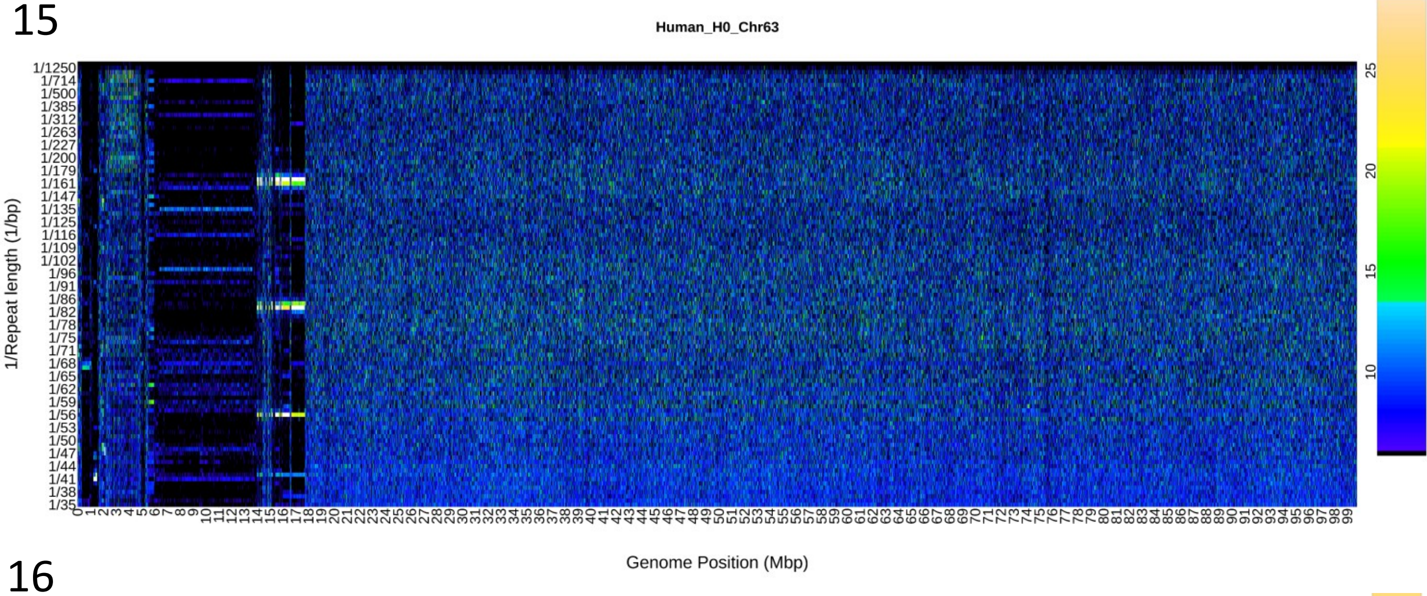

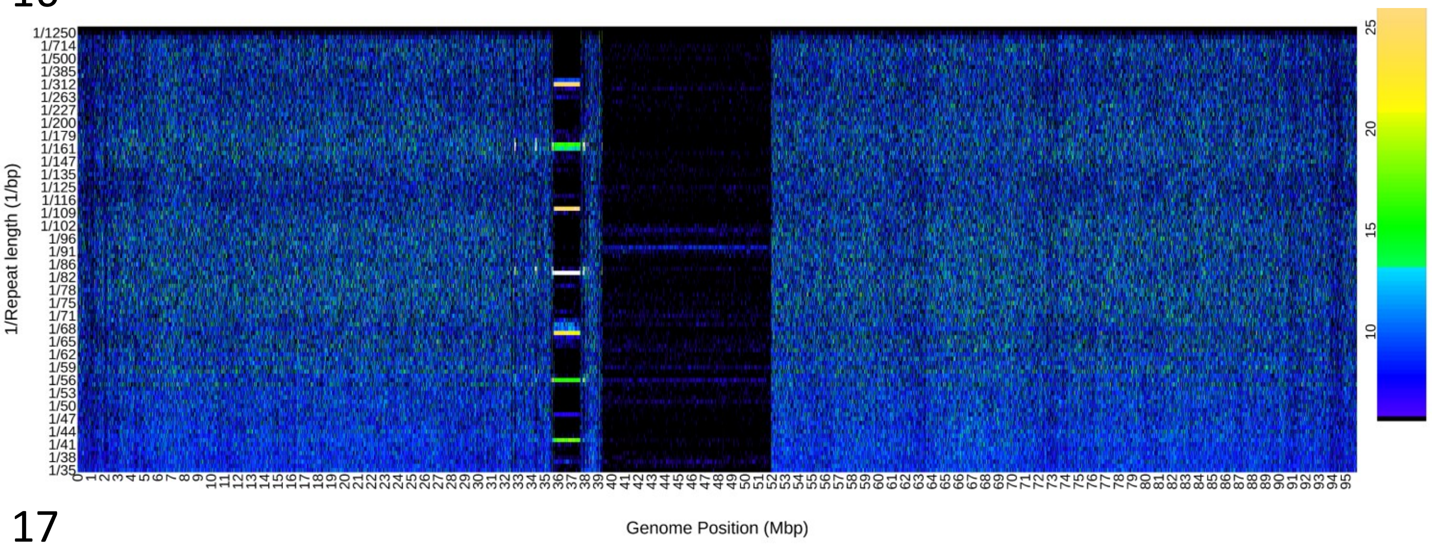

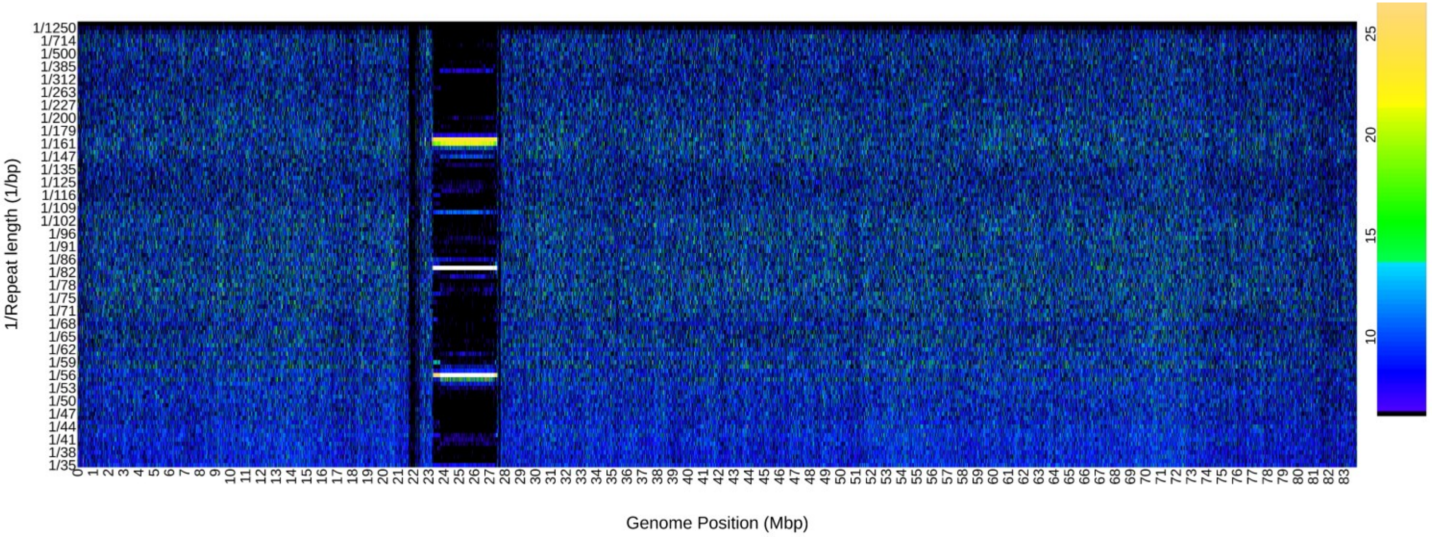

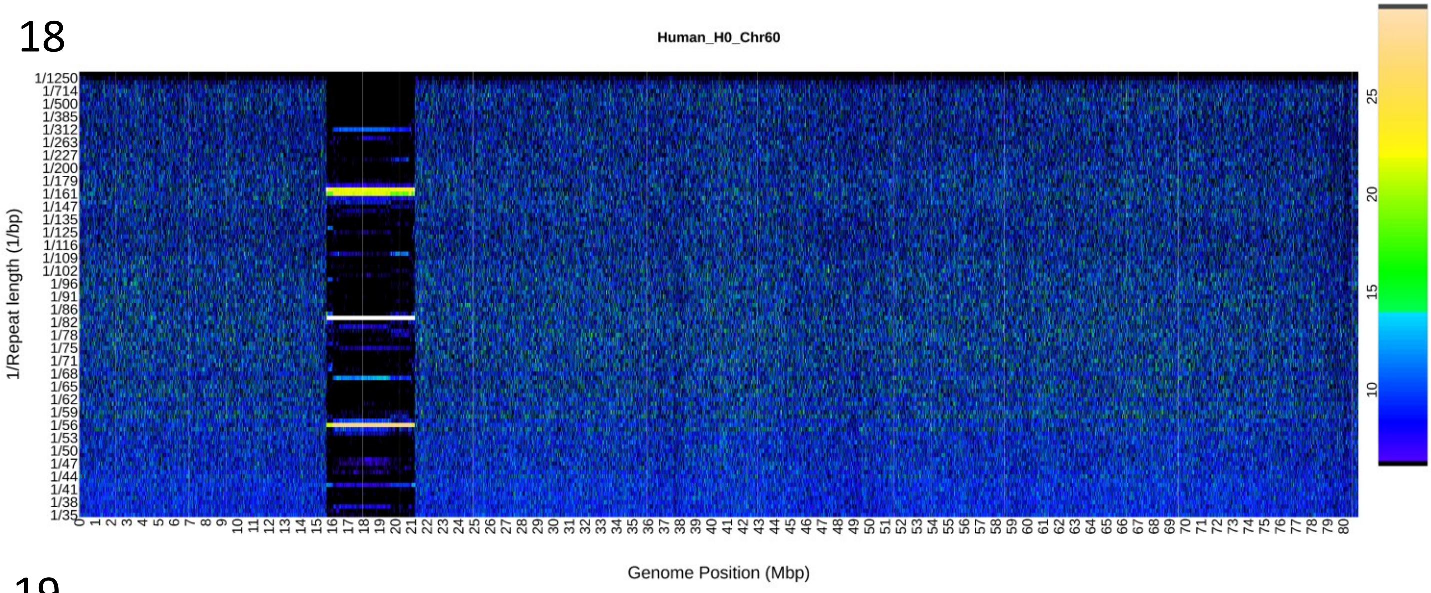

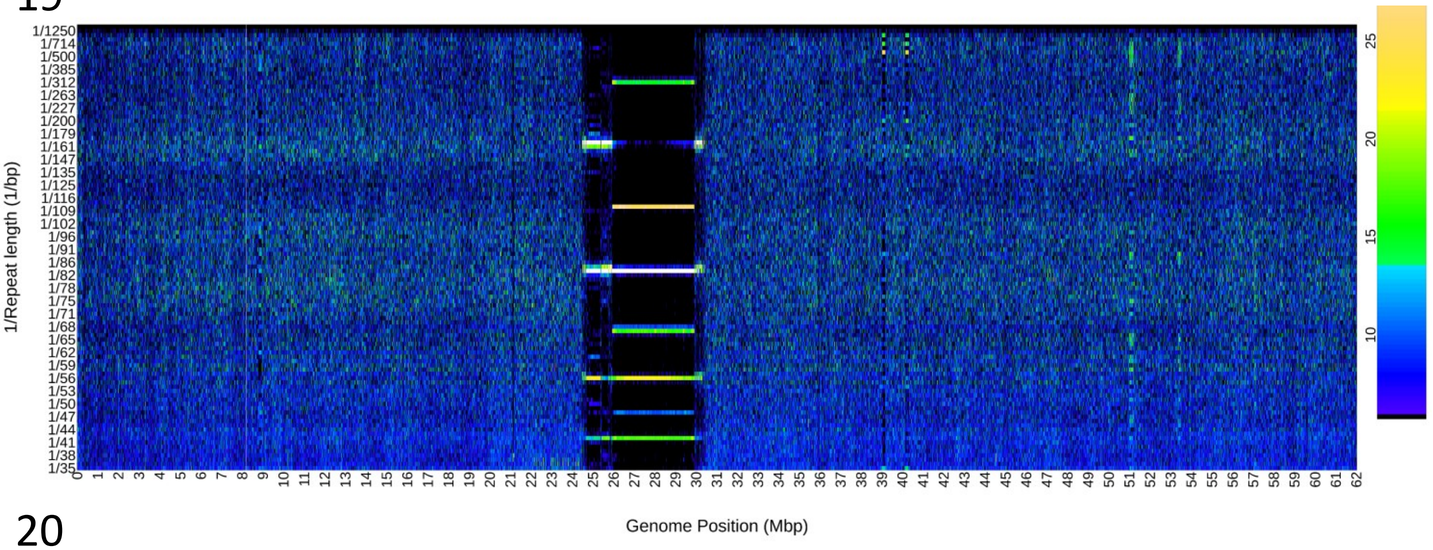

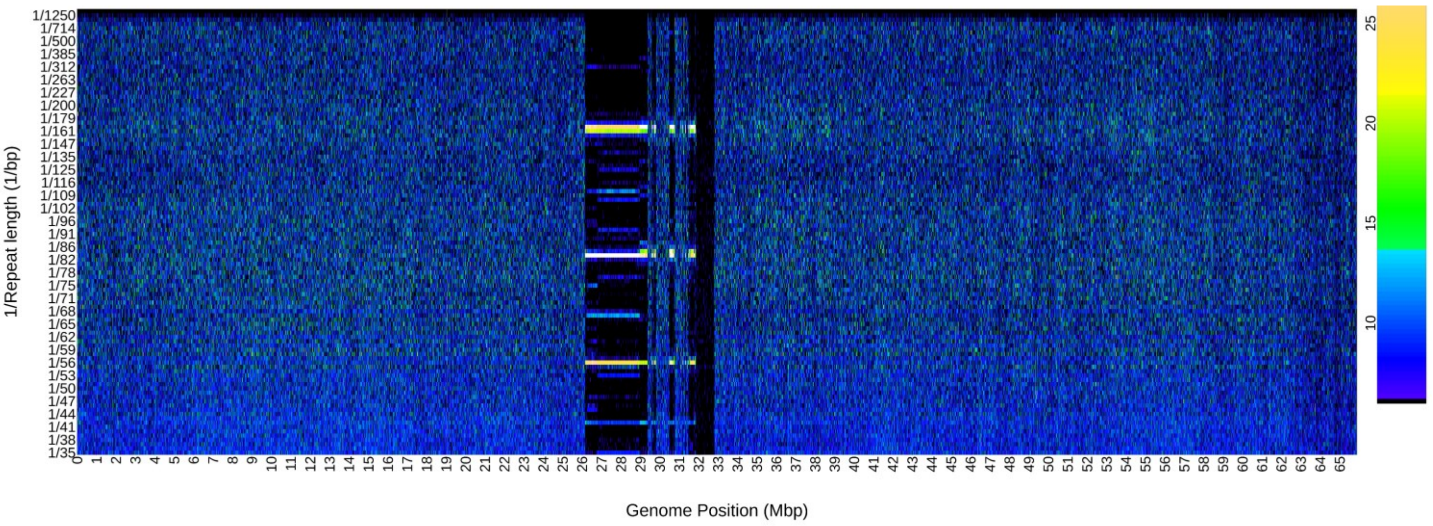

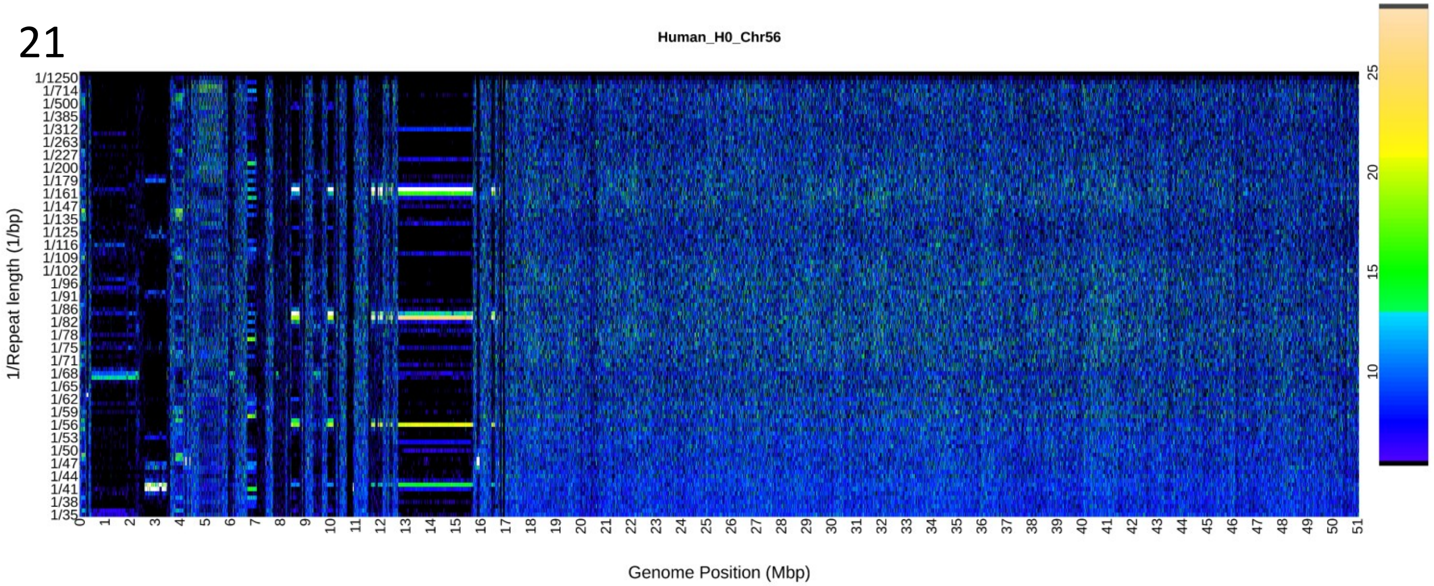
